# Increased Synapse Elimination by Inflammatory Cells Contributes to Long-lasting Post-Stroke Memory Dysfunction in Old Mice

**DOI:** 10.1101/2025.06.13.659591

**Authors:** Zahra Shabani, Peipei Pan, Qifeng Li, Zhanqiang Wang, Kun Leng, Shalika Sangras, Kang Huo, Alka Yadav, Calvin Wang, Joshua Shi, Gregory Chinn, Nikolas Kyritsis, Adam R. Ferguson, Judith Hellman, Hua Su

**Affiliations:** Center for Cerebrovascular Research, University of California, San Francisco; Department of Anesthesia and Perioperative Care, University of California, San Francisco; Department of Neurological Surgery, University of California, San Francisco

**Keywords:** Stroke, Aging, Memory dysfunction, Synapse elimination, Neuroinflammation

## Abstract

Old patients are more likely to experience memory dysfunction than young patients after a stroke. It has been reported that brain astrocytes and microglia cause excessive removal of synapses at the acute and subacute stages of stroke, and inhibition of their phagocytosis improved neurobehavioral outcomes. We hypothesized that memory dysfunction in old subjects is associated with increased synapse removal by inflammatory cells. Ischemic stroke was induced in young (2-month-old) and old (15-18-month-old) mice. Memory functions were analyzed by the Y-maze test weekly for 8 weeks and the novel object recognition (NOR) test at 7 days before and 8 weeks post-stroke. We have also created a tibia fracture 6 hours before stroke injury in young mice, to test if the activation of α7-nicotinic acetylcholine receptor (nAchRs) reduces inflammatory cells and synapse elimination. Brains were collected 8 weeks after the induction of ischemic stroke. Transcriptome changes, neuronal injuries, neuroinflammation, synapse removal, and neurite outgrowth were analyzed. We found that old mice developed long-term memory dysfunction after ischemic stroke, which was not seen in young mice. Old mice showed larger infarct volume, higher neuroinflammation, and more synapses engulfed by microglia/macrophages and astrocytes in the peri-atrophic region and hippocampi than young mice. More synapse-engulfing astrocytes than microglia/macrophages were present in the peri-atrophic region and the ipsilateral hippocampi, suggesting that reactive astrocytes contributed more than activated microglia/macrophages in synapse removal. Activation of α7-nAchRs in mice subjected to tibia fracture 6 hours before ischemic injury reduced synapse removal by microglia/macrophages and astrocytes in the hippocampi. Our study indicated that an increase in synaptic elements by inflammatory cells contributes to the long-lasting memory deficit after stroke in old mice. Astrocytes may contribute more than microglia/macrophages in synapse removal. Inhibition of neuroinflammation by activating α7-nAchRs can reduce synapse loss and thus may improve post-stroke memory function.

## Introduction

Stroke is the second leading cause of death and disability worldwide, and the fifth leading cause of death in the United States [1, 2]. Acute ischemic stroke accounts for 85% of the annual 600,000 strokes in the United States, with a 20% to 50% mortality rate [3]. Approximately 75–89% of strokes occur in people over the age of 65 years, and the stroke rate doubles for each decade after the age of 55 years. Old patients show higher mortality, severe neurological deficits, and slower recovery than young patients [4]. Age is one of the strongest non-modifiable risk factors for ischemic stroke, which is correlated with higher stroke prevalence and incidence, as well as slower post-stroke functional recovery in both human stroke patients and experimental models, leading to reduced life span and elevated cost of health care [5–8]. As the US’s aging demographics increase, memory dysfunction becomes a prominent health problem and a significant economic burden for society.

Although acute deprivation of oxygen and nutrients results in primary cell death, inflammation causes secondary cell death and plays an essential role in the development of neurological deficits in stroke patients [9]. Up to 80% of stroke survivors suffer from cognitive deficits [10]. Profound inflammatory response induced by ischemic stroke plays an important role in stroke outcome [11]. Doyle et al. demonstrated that delayed cognitive impairment in patients and mice following stroke is associated with a B-lymphocyte response [12]. A recent study suggested that post-stroke neuroinflammation is a diffuse process, which is more prominent in the ipsilateral hemisphere and minor in the contralateral hemisphere, and contributes to post-stroke cognitive impairment presentations [13].

Following a stroke, patients develop deficits in various cognitive domains. Memory impairment affects around 25% to 30% of patients in the acute phase and 9% to 15% in the chronic phase (>6 months) [11]. However, the association of neuroinflammation and cognitive dysfunction is not clear. Activated inflammatory cells can potentially eliminate synapses [14], leading to white matter damage and cognitive dysfunction [15, 16]. Neuroinflammatory response can abrogate memory-enhancing synaptic plasticity [17].

Microglia and astrocytes are highly responsive resident brain cells activated in response to stroke injuries. Astrocytic processes are dynamically associated with synapses, functionally interact with dendritic spines and synaptic terminals. These astrocytic contacts regulate the maturation of dendritic spines and their plastic structural remodeling. Microglia do the same [18]. It has been shown that reactive microgliosis and astrogliosis play important roles in mediating synapse engulfment in pathologically distinct murine stroke models. Both microglia/macrophages and astrocytes can eliminate synapses in the ischemic stroke brain at acute and subacute stages. In hemorrhagic stroke, microglia/macrophages but not astrocytes play a major role in removing synapses [19].

Transcriptomic profiling showed that both microglia and astrocytes were enriched in synapse engulfment pathway-related genes, including phagocytic receptors, and intracellular molecules [19, 20]. During aging, microglia undergo marked phenotypic and functional changes, including increased cell numbers, dystrophic morphology, impaired phagocytosis, reduced motility, and exaggerated response to inflammatory stimuli. Previous studies have highlighted the contribution of microglia/macrophages and astrocytes to neuronal debris clearance during the early stage of stroke. It is unclear whether they are still phagocytic and engulf synapses at the subacute and recovery stages [19].

We previously found that young adult mice developed long-lasting memory dysfunction when a tibia fracture preceded ischemic stroke by 6 hours, coinciding with an accumulation of CX3C chemokine receptor 1^+^ (Cx3cr1^+^) and CD68^+^ cells [21], impairment of the blood-brain barrier (BBB) and reduction of pericytes in the hippocampus, and damage of the striatum white matter [22], in contrast to the temporary memory dysfunction (<1 week) caused by BF alone [23, 24]. These data suggest that pericyte reduction and BBB impairment augment inflammatory responses in the hippocampus, which may lead to observed neuronal damage and memory dysfunction in mice with bone fractures and stroke.

Infarctions anywhere in the brain can induce widespread disruption of functional networks of the cortical regions. The hippocampus is an important brain structure for episodic memory formation, encoding, and long-term consolidation of new information [25]. Several studies reported that astrocytes and microglia/macrophages in various regions of the human brain express α7-nicotinic acetylcholine receptor (α7nAChRs), which mediate cholinergic anti-inflammatory effects [26, 27]. A better understanding of the neural mechanisms that underlie the development of persistent memory impairment following stroke is important for both early diagnosis and the development of targeted treatments [10]. In this study, we tested whether ischemic stroke results in long-term memory dysfunction and more severe neuroinflammation in old mice than young mice. We also analyzed whether activated microglia/macrophages and reactive astrocytes engulf synapses at the later stage of ischemic stroke, and which cell plays the major role. We found that old mice develop memory dysfunction that lasts 8 weeks post-ischemic stroke when the study was stopped. More activated microglia/macrophages and reactive astrocytes were present in the peri-atrophic region and hippocampi of old mice than young mice, suggesting more severe neuroinflammation. Both microglia/macrophages and astrocytes were engulfing synapses. There were more astrocytes and synapse-engulfing astrocytes than microglia/macrophages and synapse-engulfing microglia/macrophages in the hippocampi of old mice, suggesting that astrocytes play a major role in eliminating synapses in the hippocampi of old mice at the later stage of ischemic stroke. In addition, we have found that activation of nAchRs reduced synapse engulfing microglia/macrophages and astrocytes in the hippocampi of young mice with tibia fracture 6 hours before stroke Injury, suggesting that reduction of synapse engulfing by inflammatory cells contribute to the improvement of memory function by activating nAchRs in young mice with double injuries that we had observed before [28].

## Materials and Methods

### Animals

Young (2-month-old) and old (15-18-month-old) wild-type C57BL/6 mice (The Jackson Laboratory, Sacramento, CA) were housed in the pathogen-free, air-conditioned animal facility (24±1°C; 12 hours light/dark cycles) with free access to standard chow and water ad libitum at the Zuckerberg San Francisco General Hospital. Nearly equal numbers of female and male mice were used in both young and old groups. Young and old mice were assigned to different groups randomly. All experiments were approved by the Institutional Animal Care and Use Committee at the University of California, San Francisco, and conformed to National Institutes of Health guidelines for the care and use of laboratory animals.

### Tibia fracture surgery

The tibia fracture was done as described in our previous papers [21, 28, 29] in an aseptic condition. Mice were anesthetized by 4% isoflurane inhalation until no pinch-paw reflex was observed. The anesthesia was maintained during the surgical procedures by providing 1-2% isoflurane. A longitudinal incision was made from the knee to the midshaft of the tibia. The subcutaneous tissues and muscles were pushed aside to expose the patellar tendon and the tibia periosteum. A 0.5 mm hole in the proximal tibia was drilled with a 25-gauge needle just beneath and medial to the patellar tendon to enter the intramedullary canal. A 0.38 mm stainless steel rod was inserted in the canal until resistance was felt. The fibula and the muscles surrounding the tibia were isolated, and the periosteum was stripped over a distance of 10 mm circumferentially. An osteotomy was performed with scissors at the junction of the middle and distal third of the tibia. The wound was closed. The mice were allowed to recover spontaneously from anesthesia in warmed cages.

### Permanent middle cerebral artery occlusion (pMCAO) procedure

The pMCAO surgeries were performed as previously described [28, 30]. Briefly, mice were anesthetized as described above. A small vertical incision (about 1cm long) was made between the left orbit and tragus to expose the temporal bone. A small piece of skull/dura (approximately 2 mm^2^) was drilled and carefully removed to expose the distal end of the middle cerebral artery (MCA). The MCA was gently coagulated using a bipolar coagulating forceps (Codman & Shurtleff, Inc. Raynham, MA), and saline was applied during the coagulation to prevent heat injury. The skin was sutured. The mice were allowed to recover in a heated chamber for about 30 min. The sham surgery procedure was identical to pMCAO procedure except that no coagulation of MCA. The body weight was measured right before the surgery and weekly for 8 weeks after the surgery. The blood pressure was measured using CODA Non-Invasive Blood Pressure System (Kent Scientific Corporation, Torrington, CT) at four different time points: before surgery (baseline value), right after anesthesia induction, 10 minutes after surgery, and recovery from anesthesia (24 hours after surgery). The study designs are shown in **Supplementary Fig. 1**.

The old mice had higher blood pressure at baseline (p<0.001) and 24 hours after surgery (recovery stage, p=0.001) than young mice (**Supplementary Fig. 2**), and a higher body weight throughout the experimental period than young mice (**Supplementary Fig. 3**). No significant body weight change was detected in young and old mice during the experimental period (**Supplementary Fig. 3**).

### PHA568487 (PHA) treatment

PHA (Tocris Bioscience, Bristol, UK), a selective agonist of α7 nAchR, was diluted in 0.9% saline before use and was injected intraperitoneally (0.8 mg/kg) immediately before tibia fracture and one day after pMCAO, as we have done previously [28]. Saline was injected similarly to the control mice. The treatment scheme is shown in Supplementary Fig. 1b.

### Y-maze test and novel object recognition (NOR) test

The Y-maze test was performed every week up to eight weeks after pMCAO to assess the spatial working memory of mice, as we have done in our previous studies (**Supplementary Fig. 1a**) [21, 28]. Mice were placed in the center of a Y-shaped maze (Stroelting, Chicago, IL), consisting of three identical opaque plastic arms (8cm high), oriented 120 degrees apart and joined at a central point. Animals were allowed to explore the arms freely for 10 minutes.

Normal mice typically prefer to visit a new arm of the maze rather than returning to a previously visited one. The number and the sequence of arm entries were video recorded throughout the 10-minute test session. The percentage of spontaneous alternation was used to evaluate the mouse’s spatial memory function [31, 32].

The NOR test comprises three phases: the habituation phase (days 1-2), the familiarization phase (day 3), and the testing phase (day 4). During the habituation phase, mice were allowed to acclimate to a clear plastic open-field arena (45 cm x 24 cm x 20 cm) for 10 minutes per day. During the familiarization phase, each mouse was allowed to explore two identical objects in the arena for 5 minutes. During the testing phase, each mouse was allowed to explore one familiar object that was encountered in the familiarization phase and one novel object for 5 minutes.

The testing phase was recorded by a night vision camera (ELP1 Megapixel Day Night Vision, China), and the exploratory behavior was analyzed using the SMART 3.0 video tracking software (Panlab, Spain). Normal mice spend more time exploring the novel object than the familiar object [33]. The preferential exploration towards the novel object is calculated by dividing the time spent exploring the novel object by the total exploration time for both objects throughout the first 20 seconds of object interaction. In this study, the NOR tests were done 7 days before and 8 weeks after the pMCAO, as we have done before (**Supplementary Fig. 1a**) [21, 28].

### Tissue Collection

The mice were sacrificed at 8 weeks poststroke. For immunostaining, the brains were removed, frozen on dry ice immediately, and then stored in a -80°C freezer until cryosection. Brain samples were cut into 20-μm-thick coronal sections using a cryostat (Leica Microsystems, Wetzlar, Germany). The sections collected from the peri-atrophic region (from bregma -1.2 mm to bregma -1.4 mm) and hippocampal region (from bregma -1.6 mm to bregma -2.3 mm) were used for histological and immunohistochemical stains. For RNA sequencing and western blot analyses, the brain was rapidly removed, and the cortical and hippocampal regions from both hemispheres were dissected on a metal plate on top of ice, frozen on dry ice immediately, and then stored in -80°C until use.

### Measurement of atrophy volume

Cresyl violet staining was carried out by sampling 10 cortical sections/mouse, spaced 200 μm apart. The hemisphere areas were outlined and quantified using Photoshop. The atrophic area of each brain section was calculated by subtracting the area of the ipsilateral hemisphere from the area of the contralateral hemisphere. The atrophic volume was calculated by multiplying the sum area of the atrophic regions of 10 brain sections by 200 μm.

### Immunostaining

Brain sections were fixed in methanol for 20 min at -20°C and rinsed with PBS three times. For CD68/synaptophysin and GFAP/synaptophysin co-staining, brain sections were blocked with 2.5% normal donkey serum for 1 hour at room temperature and then incubated with rat anti-CD68 (1:300, MCA1957, Bio-Rad, Hercules, CA), or rat anti-GFAP (1:400, 13-0300, Thermo Fisher Scientific, Waltham, MA) together with rabbit anti-synaptophysin (1:200, Cat. ab52636, Abcam, Cambridge, UK) antibodies at 4°C for overnight. On the following day, after rinsing with PBS three times, the brain sections were incubated with secondary antibodies: Alexa Fluor 488-conjugated donkey anti-rat antibody (1:200, Cat. A 21208, Invitrogen, Waltham, MA), or Alexa Fluor 555-conjugated donkey anti-rabbit antibody (1:300, Invitrogen) for 1 hour at room temperature. After rinsing with PBS three times, the brain sections were mounted using Vectashield antifade mounting media containing DAPI (Vector Laboratories, Inc., Newark, CA). Images were acquired using a fluorescence microscope (Boprevo BZ-9000, Keyence, Osaka, Japan).

### Western blot

Protein were extracted from brain tissues (1 mm^3^) of the atrophic region, corresponding contralateral cortex, hippocampus ipsilateral and contralateral to stroke injury using a cell lysis buffer (Cell Signaling Thchnology, Danvers, MA) supplemented with 1 mM PMSF (Cell Signaling) and quantified by the Bradford method (Bio-Rad, Hercules, CA) or the BCA method (Thermo Scientific, Emeryville, CA) using a microplate reader (Emax, Molecular Devices, Sunnyvale, CA). Protein samples were run into 4–20% Tris-Glycine gels (Bio-Rad, Hercules, CA) and transferred onto PVDF membranes (Bio-Rad). Immunoblotting was performed using primary antibodies specific to GFAP (rat anti-GFAP, 1:500, 13-0300, Invitrogen) and beta actin (mouse anti-actin, 1:2000, Cell Signaling Technology). A goat anti-rat IgG antibody (1:10,000 IRDye680CW-conjugated, 926-68071, Li-Cor, Lincoln, NE) and a goat anti-mouse IgG antibody (1:10,000, IRDye680RD-conjugated, 926-68072, Li-Cor) were used as the secondary antibody. GFAP and beta actin (control for protein loading) were detected by a Li-Cor Quantitative western blot scanner and quantified using Li-Cor imaging software (Li-Cor). The level of GFAP was normalized by the level of beta-actin.

### RNA Sequencing

The total RNAs were isolated from 1 mm^3^ of atrophic and corresponding contralateral cortical regions, and hippocampi ipsilateral and contralateral to stroke injury. RNA samples were sent to Novogene Co. (Sacramento, CA) for sequencing using the company’s standard protocol (Supplementary File S1). The outcome data were analyzed by Novogene Co.

Differential expression analysis was used to compare differences in gene expression across the experimental groups and to detect gene expression changes predominantly seen in specific experimental groups. For example, genes primarily affected by stroke were distinguished from genes that were primarily affected by age. Following differential analysis, Gene Ontology (GO) and Kyoto Encyclopedia of Genes and Genomes (KEGG) analyses were performed to detect the molecular and signaling pathway changes among groups.

We then used Weighted Gene Co-Expression Network Analysis (WGCNA), a method that identifies clusters (or “modules”) of highly co-expressed genes to uncover networks of genes that might show age-related and hemisphere-specific responses to ischemic injury. WGCNA works by grouping genes with similar expression patterns across samples. Genes involved in related biological processes tend to be co-expressed. Each module is summarized by an “eigengene,” a synthetic variable representing the average expression profile of genes in that module, allowing us to compare module activity across different experimental conditions.

### Golgi-Cox staining

Golgi-Cox staining was performed to visualize dendritic spine structure in superficial and deep cortical layer neurons and hippocampal neurons using the FD Rapid GolgiStain Kit (FD Neurotechnologies, Inc., Ellicott City, MD) according to the protocol provided by the manufacturer. In brief, mice were deeply anesthetized and intracardially perfused with saline. The whole brain was immersed in the A + B solution supplied by the kit for 15 days in the dark at room temperature, and then transferred into the C solution provided by the kit for 72 h in the dark at room temperature. Coronal sections (100 µm thick) were cut and stained according to the FD Rapid GolgiStain protocol. The images were captured by (Boprevo BZ-9000, Keyence, Osaka, Japan). Sholl analysis was conducted by counting dendritic intersections at 20 uM radial intervals from the neuronal soma using the Simple Neurite Tracer (SNT) plugin in Fiji [34]. Spine density was counted manually using apical main dendrites in the peri-atrophic region and hippocampal CA1, CA2, CA3, and DG regions. Three different neurons per mouse (a total of 9 dendrites per group) in each of these regions were used. Spine density was reported as the number of spines per 10 μm dendrite length.

### Statistical analysis

All quantifications were performed by 2-3 researchers who were blinded to the group assignment. All data were expressed as mean ± SD. Statistical analyses were performed with GraphPad Prism 9.0. Student t-test was used to compare the differences between the 2 groups. One-way ANOVA, followed by Tukey post hoc test, was used to compare differences among multiple groups; 2-way ANOVA, followed by Tukey post hoc test, was used to compare the changes according to the different levels of multiple categorical variables, such as for the Y-maze study. Statistical significance was defined as p < 0.05. Sample sizes were indicated in the figure legends.

## Results

### Old mice exhibited long-lasting memory dysfunction after stroke injuries

Data obtained from Y-maze tests weekly, and a novel object recognition (NOR) test at 7 days before and 8 weeks after the induction of the ischemic stroke were used to evaluate the impact of ischemic stroke on mouse memory function. Y-maze test showed that both young and old mice displayed 60-70% spontaneous alternation before stroke surgery, indicating that young and old mice performed equally in the test (**Fig. 1a**). However, old mice made fewer spontaneous alternations from 3 to 8 weeks after stroke than young and sham-operated old mice (p=0.013 to <0.001, **Fig. 1a)**, suggesting that old mice developed long-lasting spatial memory dysfunction after ischemic stroke. In addition, old mice made fewer entries in 10 minutes before and after ischemic stroke than young stroke mice (**Supplementary Fig. 4a**). No gender differences were found in young and old mice (**Supplementary Fig. 4b & c**).

**Fig. 1.**
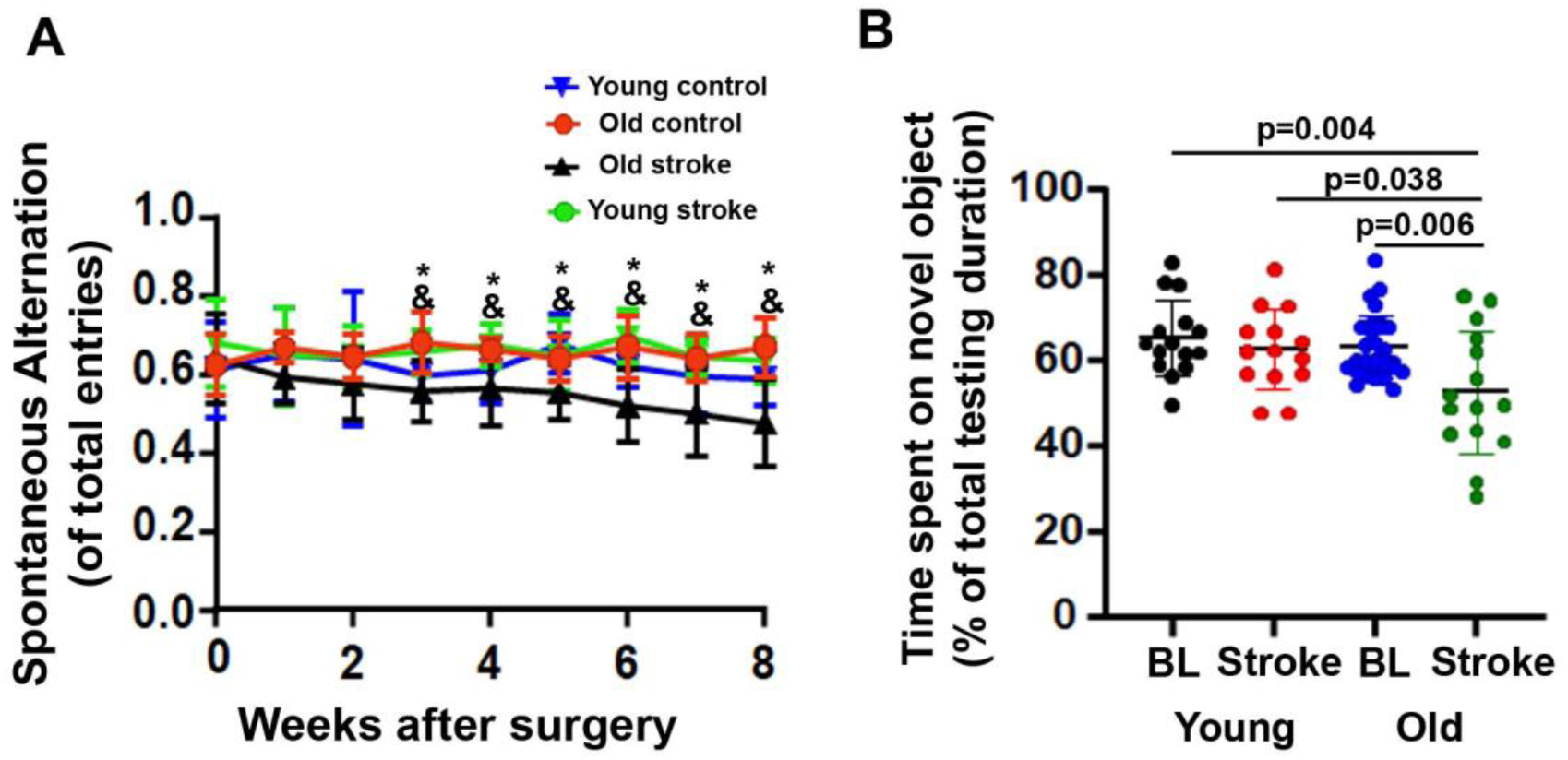
Old stroke mice showed memory dysfunction in Y-maze and NOR tests **(a).** Y-maze test showed that the old mice made fewer spontaneous alternations from 3 to 8 weeks after stroke than young stroke mice (&) and sham-operated old mice (old control, *). n=7-16. **(b).** NOR test showed that old stroke mice spent a shorter time on novel objectives than young stroke mice and young and old mice at baseline before stroke. BL. baseline. N=15-30.

For the NOR test, the baseline performance was evaluated one week before stroke injury, there was no difference between young and old mice. However, 8 weeks after stroke injury, old mice were not able to differentiate novel from familiar objects and thus spent a shorter time exploring novel objects than their baseline level (p=0.006), baseline of young mice (p=0.004), and young stroke mice (p=0.038). Young stroke mice spent similar time on novel objects after stroke as they did before stroke (Fig. 1b). There is no difference in distance traveled and average velocity between young and old mice at baseline and after stroke (**Supplementary Fig. 5**)

Overall, the results of the Y-maze and NOR tests indicate that age could be a risk factor for post-stroke memory dysfunction.

Similar to what we have reported on the acute stage of stroke [30], old mice have larger atrophy volume (p = 0.004, **Supplementary Fig. 6**), and more CD68^+^ cells in the peri-atrophic (p<0.001) and hippocampal regions ipsilateral to stroke injuries than young stroke mice (p=0.009, **Supplementary Fig. 7**) as well as more CD68^+^ cells in the ipsilateral hippocampi than the contralateral hippocampi (p<0.001, **Supplementary Fig. 7**). Old mice also had higher baseline neuroinflammation. There were more CD68^+^ cells in old brains than young brains before stroke (p=0.030, **Supplementary Fig. 8**), which may contribute to the heightened post-stroke inflammatory response.

There were also more GFAP^+^ astrocytes in the peri-atrophic region of old stroke mice than young stroke mice (p=0.049, **Supplementary Fig. 9a & c**); more GFAP^+^ astrocytes in the ipsilateral hippocampi of old stroke mice than young stroke mice (p =0.037, **Supplementary Fig. 9b & d**). However, the number of GFAP^+^ astrocytes in the ipsilateral and contralateral sides of young and old stroke mice was similar. Consistent with immunohistological analyses, western blot analyses showed that the GFAP protein level in the atrophic regions of old mice was higher than that in the atrophic regions of young stroke mice (p<0.001), and their contralateral uninjured cortex (p=0.001, **Supplementary Fig. 9e & f**). GFAP levels were similar in the hippocampi of young and old stroke mice, on the ipsilateral and contralateral sides of stroke injury (**Supplementary Fig. 9e & g**). Together, these data show that prolonged accumulation of activated microglia/macrophages and reactive astrocytes in the peri-atrophy areas and hippocampi is associated with long-term post-stroke memory decline in old mice.

### Old mice have more CD68^+^/SYP^+^ and GFAP^+^/SYP^+^ cells in the petri-atrophic and hippocampal regions after stroke

Microglia and astrocytes cause excessive removal of synapses at the early and subacute stages of stroke [19]. To examine if activated microglia/macrophages (CD86^+^) and reactive astrocytes also engulf synapses at the chronic stage of stroke, CD68^+^/SYP^+^ and GFAP^+^/SYP^+^ cells were quantified in regions illustrated in **Fig. 2a & b** on immunohistochemically stained sections. We found more CD68^+^/SYP^+^ cells in the peri-atrophic region of old stroke mice than young stroke mice (p=0.045, **Fig. 2c & d**); in the ipsilateral side than the contralateral side of the hippocampi in the old stroke mice (p=0.035, **Fig. 2f & h**). The old stroke mice also had more CD68^+^/SYP^+^ cells in the ipsilateral side of the hippocampi than in young stroke mice (p=0.042, **Fig.2f & h**). The young mice also had more CD68^+^/SYP^+^ cells in the ipsilateral side than the contralateral side of the hippocampi (p=0.042, **Fig.2f & h).**

**Fig. 2.**
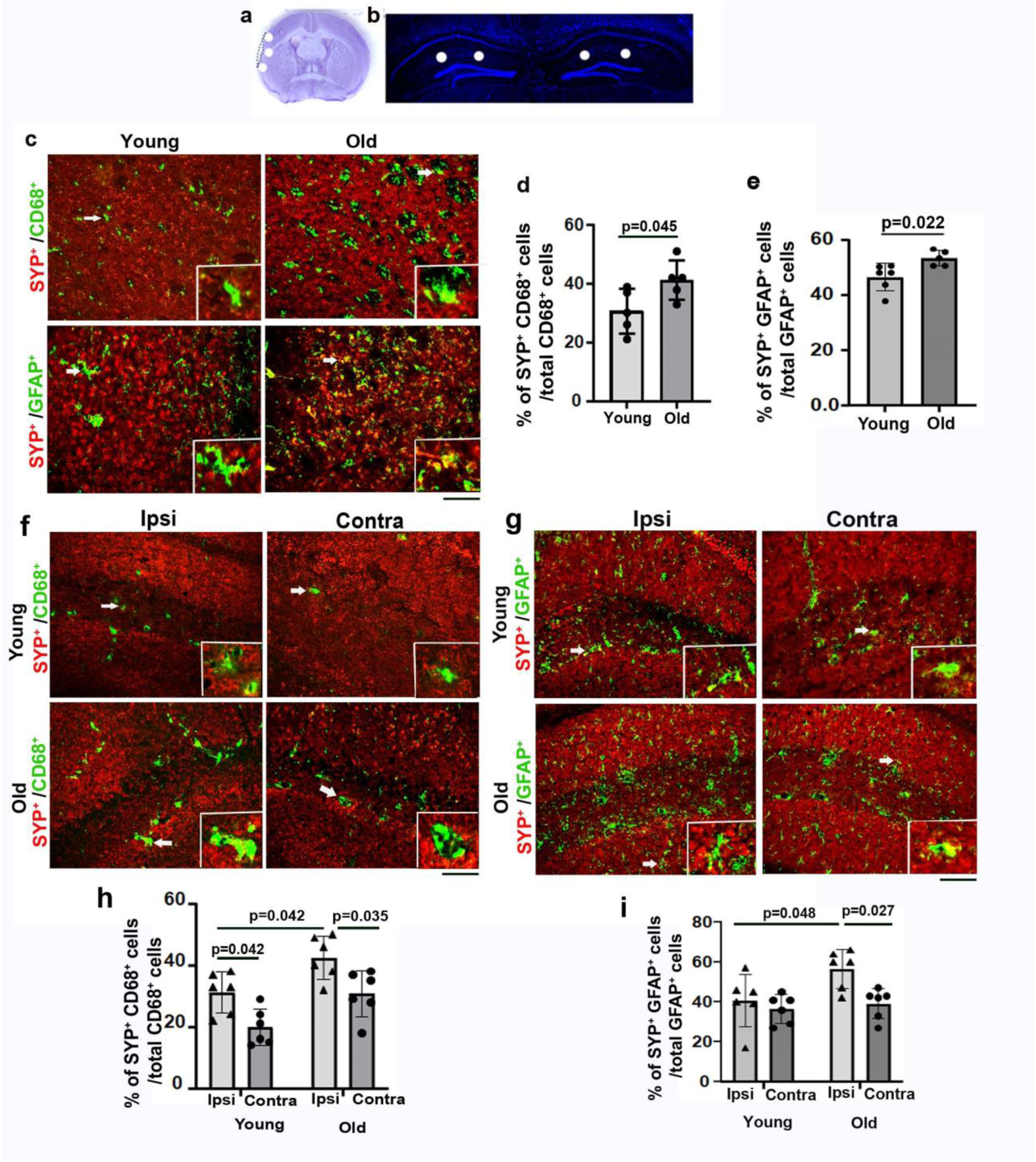
Old mice had more CD68^+^/SYN^+^ and GFAP^+^/SYN^+^ cells in the peri-atrophic regions and hippocampi ipsilateral to stroke injuries **(a).** Locations used for quantification (white spots) in peri-atrophic regions. **(b).** Locations used for quantification (white spots) hippocampal regions. **(c).** Images of peri-atrophic regions co-stained with antibodies specific to SYN (red)/CD68 (green) and SYN (red)/GFAP (green). Insertions are enlarged images of cells indicated by arrows showing colocalization of CD68/SYN and GFAP/SYN proteins. Scale bar=50µm. **(d)**. Quantification of CD68^+^/SYN^+^ among total CD68^+^ cells in peri-atrophic regions. n=5. **(e).** Quantification of GFAP^+^/SYP^+^ cells among total GFAP^+^ cells in the peri-atrophic region. N=5-6**. (f).** Images of hippocampi co-stained with antibodies specific to SYN (red) and CD68 (green). Insertions are enlarged images of cells indicated by arrows showing colocalization of CD68 and SYN proteins. Scale bar=50µm. **(g).** Images of the hippocampal regions co-stained with antibodies specific to SYN (red) and GFAP (green). Insertions are enlarged images of cells indicated by arrows to show colocalization of GFAP and SYN. Scale bar=50µm. **(h).** Quantification of CD68^+^/SYN^+^ cells among total CD68^+^ cells in the hippocampal region. n=6. **(i).** Quantification of GFAP^+^/SYP^+^ cells among total GFAP^+^ cells in the hippocampal region. n=6. Ipis: ipsilateral to stroke injury; Contra: contralateral to stroke injury. Ipis: ipsilateral to stroke injury; Contra: contralateral to stroke injury.

There were more GFAP^+^/SYP^+^ cells in the peri-atrophic region of old stroke mice than young stroke mice (p =0.022, **Fig. 2. c & e**). Old stroke mice had more GFAP^+^/SYP^+^ cells in the ipsilateral than the contralateral hippocampi (p=0.027, **Fig. 2. g & i**). There were also more GFAP^+^/SYP^+^ cells in the ipsilateral side of the hippocampi of old stroke mice than in young stroke mice (p=0.048, **Fig. 2. g & i**).

Together, the above data indicate that both CD68^+^ microglia/macrophages and reactive astrocytes are engulfing more synapses in old mice in the peri-atrophic region and hippocampus ipsilateral to stroke injury than in young mice at the chronic stage of ischemic stroke, which may contribute to the long-lasting post-stroke memory dysfunction in old mice. About 10-fold more GFAP^+^ cells were detected in old hippocampi on both ipsilateral and contralateral sides (**Fig. 3**). There were more GFAP^+^/SYN^+^ cells than CD68^+^/SYP^+^ in the peri-atrophic region (p=0.006) and the ipsilateral hippocampi (p=0.030). These data indicate that reactive astrocytes contributed more than activated microglia/macrophages in synapse engulfment in old mice at the chronic stage of stroke.

**Fig. 3.**
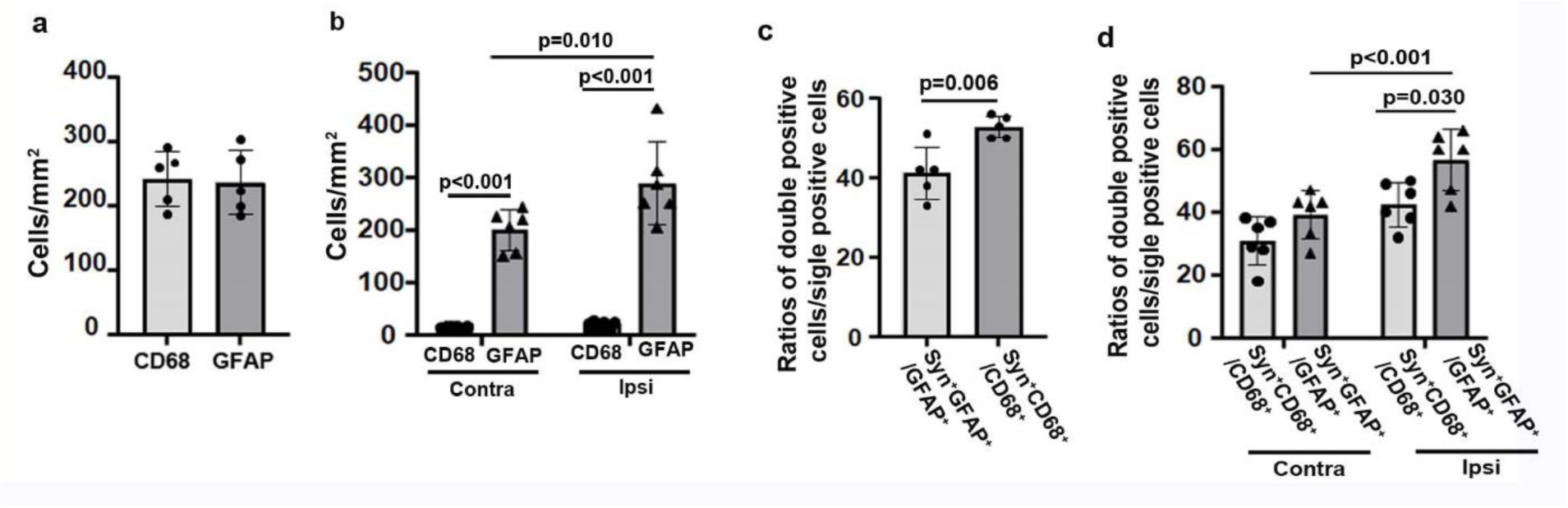
More GFAP^+^/SYN^+^ cells than CD68^+^/SYN^+^ cells in the peri-atrophic region and ipsilateral hippocampal regions of old mice **(a).** Quantification of CD68^+^ and GFAP^+^ cells in the peri-atrophic region. n=5. **(b).** Quantification of CD68^+^ and GFAP^+^ cells in the hippocampal region. n=6. **(c).** Quantification of CD68^+^/SYN^+^ and GFAP^+^/SYN^+^ in the peri-atrophic region. n=5. **(d).** Quantification of CD68^+^/SYN^+^ and GFAP^+^/SYN^+^ in the hippocampal region. n=6.

### Old mice displayed a reduced dendritic arborization of neurons in the peri-atrophic regions and ipsilateral hippocampal neurons in the CA1, CA2, CA3, and dentate gyrus (DG) regions

Golgi staining was used to track and quantify the dendritic complexities in the peri-atrophic regions indicated in **Fig. 4a** and the corresponding regions in the contralateral cortex. The average neurite length in old peri-atrophic regions was significantly shorter than old contralateral (p=0.040) and young peri-atrophic regions (p=0.016, **Fig. 4b**). There were fewer neurities in the old peri-atrophic regions than in the corresponding contralateral cortex (p=0.006) and young peri-atrophic regions (p=0.040, **Fig. 4c**). The mumber of neurities in the peri-atrophic regions of young mice were reduced compared to those in their corresponding contralateral cortex (p=0.007, **Fig. 4c**). Sholl analysis showed that there were fewer intersections in old peri-atrophic regions than in young peri-atrophic regions and old corresponding contralateral cortex (**Fig. 4d**). Area under the curve (AUCs) were significantly smaller in the old and young peri-atrophic regions compared to their corresponding contralateral areas (p<0.001, **Fig. 4e**). The spine density was reduced significantly in apical dendrites in the old peri-atrophic regions compared to the young peri-atrophic regions (p=0.032, **Fig. 4f & g**). The spine density was also lower in the peri-atrophic regions than the corresponding contralateral areas of young mice (p=0.011, **Fig. 4f & g**). The spine densities were similar in apical dendrites in old peri-atrophic regions to those in the old corresponding contralateral cortex.

**Fig. 4.**
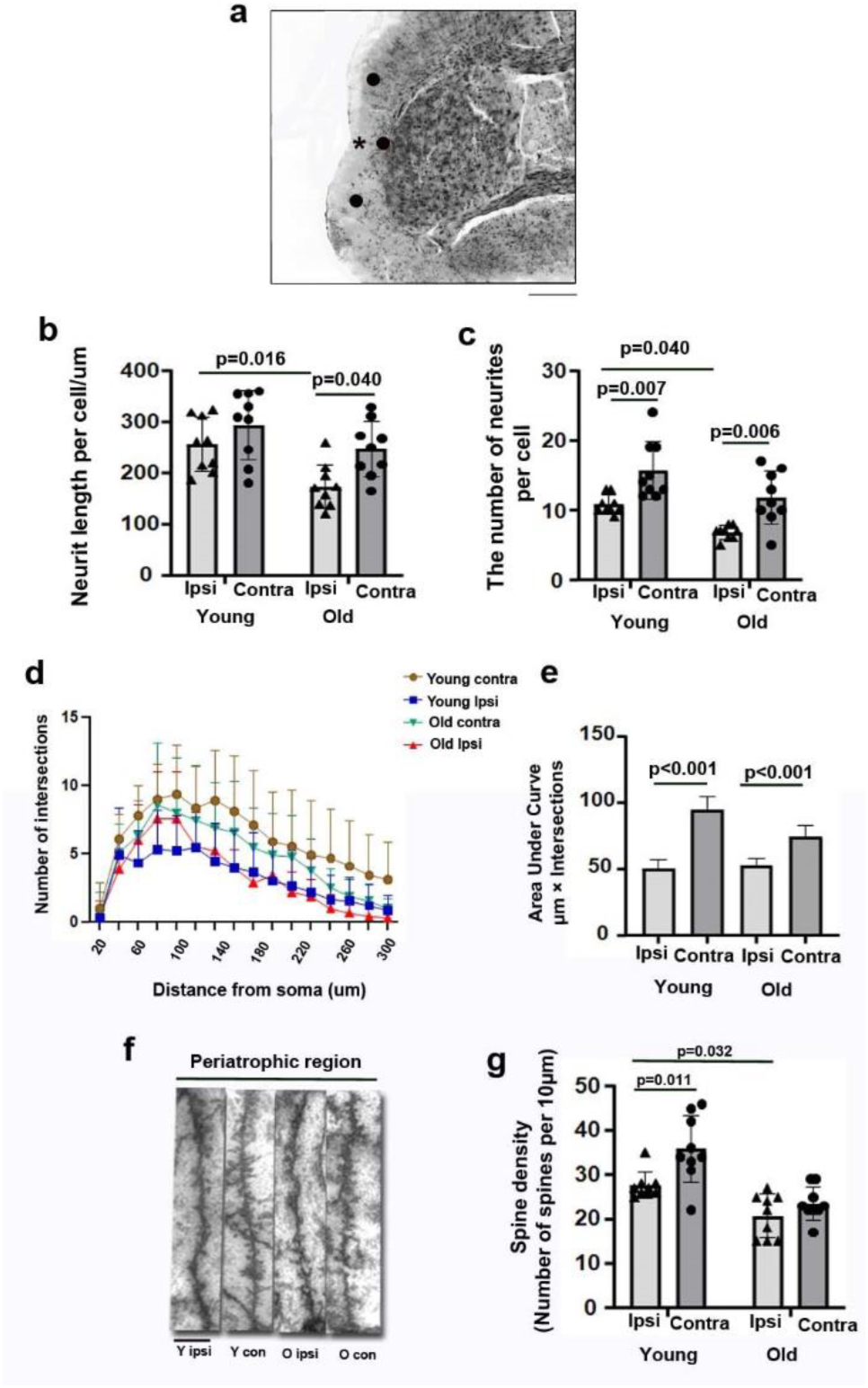
Old mice had shorter neurite length, fewer neurites, and spines in the peri-atrophic region **(a).** An image of a Golgi-stained peri-atrophic region illustrates the sites (black dots) used for analysis. *: atrophic region. Scale bar=1 mm. **(b).** Quantification of neurite length. **(c.)** Quantification of neurite number. **(d).** Quantification of intersections in Sholl analysis. **(e).** Quantification of AUC. **(f).** Representative image of selected segments of apical dendrites for spine quantification. Scale bar=5µm. **(g).** Spine density. AUCs, areas under the curves that are shown in Sholl analysis.

We have also analyzed the dendritic complexity of the neurons in the CA1, CA2, CA3, and DG (**Fig. 5a**). Apical dendrites in old ipsilateral CA1 were shorter (p=0.020, **Fig. 5b**) and have fewer neurites than those in the young ipsilateral CA1 (p=0.008, **Fig. 5c**). The total length and the number of neurites of apical dendrites were shorter and have fewer branches in the ipsilateral CA1 than that in the contralateral CA1 in both young and old stroke mice (p<0.001, **Fig. 5b & c**). Sholl analysis shows that the dendrites of neurons in the ipsilateral CA1 of old mice have shorter radial distances (fewer intersections) than those in ipsilateral CA1 of young mice and contralateral CA1 of old mice (**Fig. 5d**). The AUC was significantly smaller in the old ipsilateral CA1 than the contralateral CA1 (p=0.036, **Fig. 5e**).

**Fig. 5.**
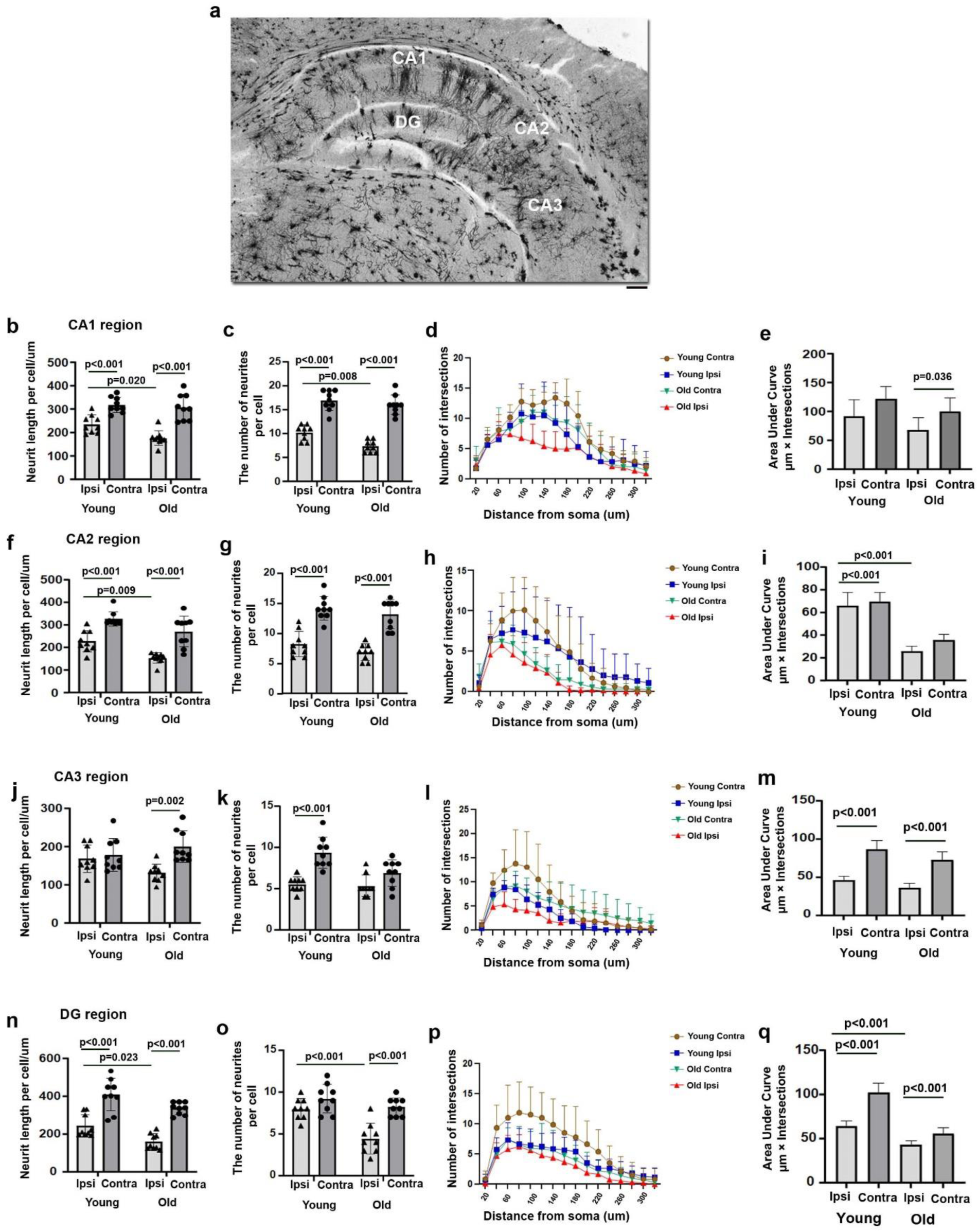
Old mice had shorter neuronal neurite length and fewer neurites in the ipsilateral CA1, CA2, CA3, and DG regions (a). An image of a Golgi-stained hippocampus illustrates the regions analyzed. Scale bar: 100 µm. (b). Quantification of neurite length in the CA1 region. (c). Quantification of neurite numbers in the CA1 region. (d). Quantification of intersections in Sholl analysis in the CA1 region. (e). Quantification of AUCs in the CA1 region. (f). Quantification of neurite length in the CA2 region. (g). Quantification of neurite number in the CA2 region. (h). Quantification of intersections in Sholl analysis in the CA2 region. (i). Quantification of AUCs in the CA2 region. (j) Quantification of neurite length in the CA3 region. (k). Quantification of neurites number in the CA3 region. (l). Quantification of intersections in Sholl analysis in the CA3 region. (m). Quantification of AUCs in the CA3 region. (n). Quantification of neurite length in the DG region. (o). Quantification of neurite number in the DG region. (p). Quantification of intersections in Sholl analysis in the DG region. (q). Quantification of AUCs in the DG region.

The dendrites of neurons in old ipsilateral CA2 have shorter total length than those in young ipsilateral CA2 region (p=0.009, **Fig. 5f**). The total length and the number of neurites of the old and young CA2 basal dendrites in ipsilateral hippocampi were significantly reduced compared to the contralateral sides (p<0.001, **Fig. 5f & g**). Sholl analysis shows that the dendrites of neurons in the ipsilateral CA2 of old mice have shorter radial distances (fewer intersections) than those in ipsilateral CA2 in young mice and contralateral CA2 of old mice (**Fig. 5h**). The AUC was significantly smaller in the old ipsilateral CA2 than in the young ipsilateral CA2 (p<0.001, **Fig. 5i**). The AUC was also smaller in the ipsilateral than contralaterial CA2 in young mice (p<0.001, **Fig. 5i**).

The total length of apical dendrites in the old ipsilateral CA3 was significantly shorter than those in the contralateral CA3 (p=0.002, **Fig. 5j**). The number of neurites of the young basal dendrites in ipsilateral CA3 was reduced compared to that in contralateral CA3 (p<0.001, **Fig. 5k**). The dendrites of neurons in old ipsilateral CA3 have shorter radial distances (fewer interactions) than those in young ipsilateral CA3 and old contralateral CA3 (**Fig. 5l**). The AUC was significantly smaller in the old and young ipsilateral CA3 than in their contralateral CA3 (p<0.001, **Fig. 5m**).

In the DG region, the total length of apical dendrites in old ipsilateral hippocampi was significantly shorter than those in young ipsilateral hippocampi (p=0.023, **Fig. 5n**). Both the neurites and the total length of the old and young DG apical dendrites in ipsilateral hippocampi were significantly reduced compared to contralateral hippocampi (p<0.001, **Fig. 5n & o**). The dendrites of neurons in old ipsilateral DG have shorter radial distances (fewer intersections) than those in young ipsilateral DG and old contralateral DG (**Fig. 5p**). The AUC was smaller in the old ipsilateral DG than young ipsilateral DG (p<0.001, **Fig. 5**q). The AUCs were also smaller in the ipsilateral DG compared to the contralateral DG in both old and young mice (p<0.001, **Fig. 5q**).

The spine densities were quantified in apical main dendrites in the CA1, CA2, CA3, and DG regions. The spine densities of apical dendrites were lower in the young (p=0.002) and old (p=0.033) ipsilateral CA1 than the contralateral CA1. (**Fig. 6a & b**). Spine density was significantly lower in apical dendrites in the old ipsilateral CA2 compared to the young ipsilateral CA2 (p=0.002, **Fig. 6c & d**). In the CA3 region, spine densities of apical dendrites in the young and old ipsilateral region were lower than the contralateral sides (p<0.001, **Fig. 6e & f**). The spine density was significantly lower in apical dendrites in the old ipsilateral DG region than in the young ipsilateral DG (p=0.018, **Fig. 6g & h**). Old ipsilateral DG region showed lower spine densities than old contralateral DG (p=0.024, **Fig. 6g & h**).

**Fig. 6.**
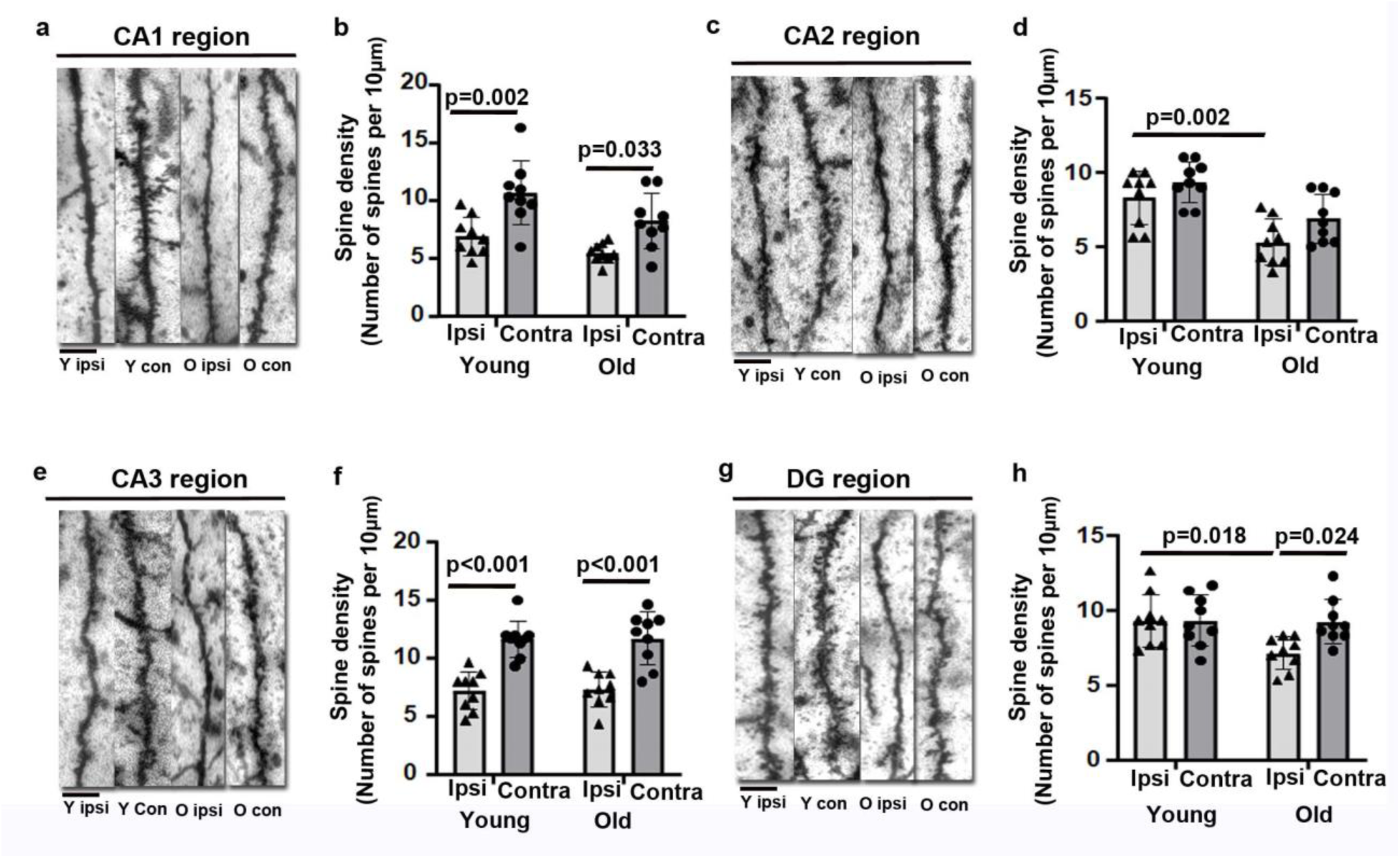
Old mice have fewer dendritic spines in ipsilateral neurons in the CA1, CA2, CA3, and DG regions **(a).** Representative images of apical dendritic segments analyzed in the CA1 region. **(b).** Quantitative spine densities of apical dendrites in the CA1. **(c).** Representative images of apical dendritic segments analyzed in the CA2. **(d).** Quantitative spine densities in apical dendrites in the CA2. **(e).** Representative images of apical dendritic segments analyzed in the CA3. **(f).** Quantitative spine densities in apical dendrites in the CA3. **(g).** Representative images of apical dendritic segments analyzed in the DG. **(h).** Quantitative spine densities in apical dendrites in the DGs. Scale bars=5 µm.

### Ischemic stroke injury induced higher neuroinflammation and oxidative stress in old mice compared to young mice

We have analyzed transcriptomes in young and old mice in uninjured and injured cortical regions and hippocampi collected 8 weeks after ischemia. Different brain regions show distinct gene expression patterns (**Fig. 7**).

**Fig. 7.**
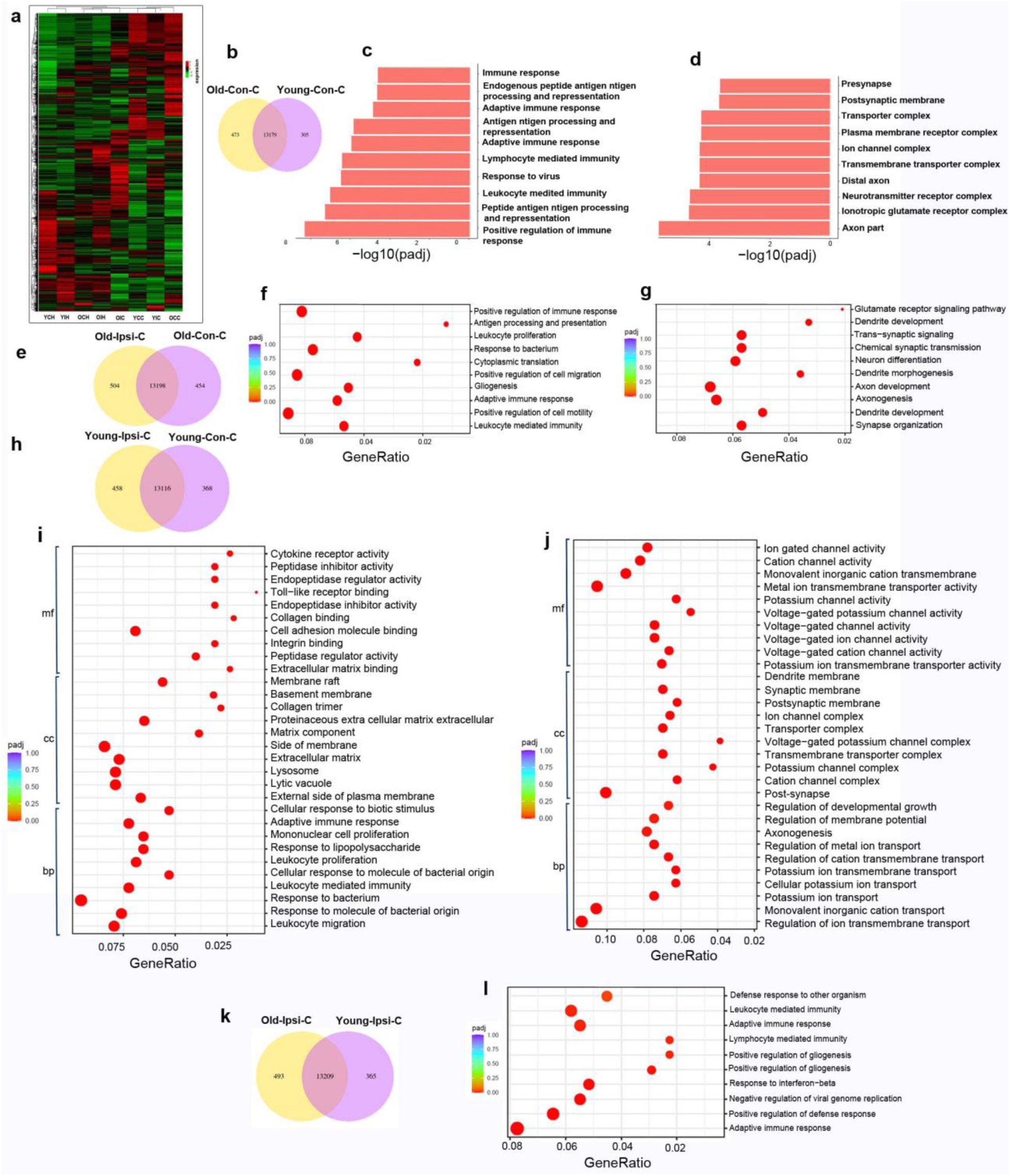
Differential gene expression and pathways changes identified by RNA-seq (a). A heatmap shows distinct gene expression patterns in different brain regions.YCH: young contralateral hippocampi; YIH: young ipsilateral hippocampi; OCH: old contralateral hippocampi; OIH: old ipsilateral hippocampi; OIC: old ipsilateral cortex; YCC: young contralateral cortex; YIC: young ipsilateral cortex; OCC: old contralateral cortex. n=3. (b). Venn diagram of differential gene expression in the uninjured old cortex compared to the uninjured young cortex. (c). The top 10 up-regulated biological pathways in the uninjured old cortex compared to the young cortex identified by GO analysis. Most of them are inflammation-related pathways. (d). The top 10 down-regulated biological pathways in the unjured old cortex identified by Go analysis compared to the young cortex. Most of them are related to cell-cell transportation and neurite outgrowth. (e). Venn diagram of differential gene expression in the atrophic region compared to the corresponding contralateral cortex of old mice. (f). GO dot plot shows the top 10 up-regulated pathways in the atrophic region compared to the corresponding contralateral cortex of old mice. Most of them are related to inflammation pathways. (g). GO dot plot shows the top 10 downregulated pathways in the atrophic region compared to the corresponding contralateral cortex of old mice. Most of them are related to synaptic transmission and neurite outgrowth. (h). Venn diagram of differential gene expression in the atrophic region compared to the corresponding contralateral cortex of young mice. (i). GO dot plot shows top 10 up-regulated pathways in biological pathways (bp), cellular component (cc), and molecular function (mf) in the atrophic region compared to the corresponding contralateral cortex of young mice. Most of them are related to inflammation-related pathways. (j). GO dot plot shows the top 10 down-regulated pathways in biological pathways (bp), cellular component (cc), and molecular function (mf) in the atrophic region compared to the corresponding contralateral cortex of young mice. Most of them are related to synaptic transmission and neurite outgrowth. (k). Venn diagram of differential gene expression in the old atrophic regions compared to the young atrophic regions. (l). Inflammation pathways were upregulated in the atrophic region of old mice compared to those in young mice (top 10 upregulated pathways identified by GO analysis). n=3.

In the uninjured cortex, 473 genes were uniqualy expression in old mice and 305 genes were uniqually expression in young cortex (**Fig.7b**). GO analysis showed that inflammation pathways were upregulated, and neurite outgrowth pathways were downregulated in the uninjured cortex of old mice compared to young mice (**Fig. 7c & d**). KEGG analysis showed that phagosome, complement and coagulation cascades, antigen processing and presentation, and glutathione metabolism were upregulated, while CAMP signaling pathway, cholinergic synapse, glutamatergic synapse, and neuroactive ligand-receptor interaction were downregulated in the uninjured cortex of the old brain compared to young brain.

Differential gene expression analyses showed that the mRNA levels of complement and coagulation cascades genes such as C3 (Adjust p=2.44X10^-8^), and C4b (Adjust p=1.25X10^-15^), inflammatory pathway genes such as GFAP (Adjust p=1.35X10^-15^) and CD68 were higher and mRNA levels of Synpo2 (Adjust p=2.66X10^-6^), Nr4a3 (Adjust p=0.0003) and Nrep (Adjust p=0.0004), which are related to the neurite outgrowth and axon regeneration, were lower in the old uninjured cortex than in young uninjured cortex. These data indicate chronic inflammation and neuronal abnormalities in the brains of old mice, which are consistent with histological analysis that old mice have more CD68^+^ than young mice in the cortex before stroke injury (**Supplementary Fig. 8**).

Differential gene expression was presented in the atrophic region and corresponding cortex of old and young mice (**Fig. e & h**). Inflammation pathways were upregulated, and synaptic transmission and neurite outgrowth were downregulated in the atrophic regions compared to the corresponding contralateral cortex in young and old mice (**Fig. 7f, g, i & j**). In contrast, synapse organization, dendrite development, axonogenesis, axon development, dendrite morphonogenesis, positive regulation of neuron differentiation, modulation of chemical synaptic transmission, regulation of dendrite development, postsynapse, synaptic membrane, postsynaptic membrane, postsynaptic specialization, asymmetric synapse, postsynaptid density, neuron to neuron synapse, transmembrane transporter complex, presynapse, transporter complex, Ion gated channel activity, and voltage-gated cation channel activity were reduced in the ipsilateral cortex of old mice compared to contralateral cortex of old mice (**Fig. 7g**).

KEGG analyses showed that phagosome, leukocyte transendothelial migration, Fc Gamma R-mediated phagocytosis, complemental coagulation cascades, cytokine-cytokine receptor interaction and antigen processing and presentation were upregulated, while dopaminergic synapse, glutamatergic synapse, axon guidance, cholinergic synapse and long-term potentiation were downregulated, in the atrophic regions compared to corresponding contralateral cortex. Go analyses showed that after ischemic stroke injury, the leukocyte-mediated immunity, lymphocyte-mediated immunity, glutathione transferase activity, glutathione peroxidase activity, antioxidant activity, peroxidases activity, oxidoreductases activity, acting on peroxide as acceptor, and glycosaminoglycan binding were increased in the ipsilateral cortex of old mice compared to young mice (**Fig. 7l**). These data suggesting that ischemic injury upregulated inflammatory and reduced neuronal function in the ipsilateral cortex of both young and old mice.

Differential gene expression analyses showed that mRNA levels of *Tlr2* (adjusted p=0.001), C3 (adjusted p=3.31X10^-10^), and GFAP (Adjusted p=1.41X10^-17^) were higher in atrophic regions than corresponding cortical regions of old mice. The mRNA levels of GFAP (Adjust p=1.41X10^-18^), CD68 (Adjust p=1.62X10^-7^), C3 (Adjust p=3.71X10^-10^), and C4b (Adjust p1.48X10^-12^) were up-regulated, and mRNA levels of Stab2 (Adjust p=3.6X10^-7^), Robo3 (Adjust p=5.91X10^-7^) were down-regulated in the old ipsilateral cortex than in the young ipsilateral cortex. Therefore, ischemic injury induced higher inflammatory response and oxidative stress and more neuronal damage in old mice than in young mice. These data are consistent with histological and western blot analysis that old mice have more CD68^+^ and GFAP^+^ cells, higher levels of GFAP protein, and increased numbers of SYN^+^ CD68^+^ and GFAP^+^ cells (**Supplementary Figs. 7-9**), and shorter neuronal neurite length, fewer neurites and dendritic spines in the peri-atrophic cortex and hippocampi ipsilateral to stroke injury than young mice (**Fig. 4**).

### Inflammation was upregulated and neurite outgrowth was downregulated in the hippocampi of old mice compared to young mice after stroke

Differential analyses showed that 1012 genes were upregulated in the hippocampi of old mice compared to young mice. Go analyses indicated that adaptive immune response, postsynapse, synaptic membrane, cation channel complex, and axon part were increased in the hippocampi contralateral to the stroke injury site in old mice compared to young mice. KEGG analyses showed that the chemokine signaling pathway, inflammatory mediator regulation of TRP channels, neurotrophin signaling pathway, and complement, Wnt signaling pathway, vascular smooth muscle contraction, GABAergic synapse, glutamatergic synapse, and cholinergic synapse, and coagulation cascade were upregulated, while gap junction, cholinergic synapse, neuroactive ligand-receptor activity, cell adhesion molecules, and axon guidance were downregulated in the ipsilateral hippocampi of old mice compared to young mice. In addition, KEGG analyses demonstrated up-regulation of complement and coagulation cascades in the hippocampi of old mice compared to young mice (Adjust p=9.49X10^-8^). Old mice showed a higher inflammatory response in the ipsilateral hippocampi in response to stroke injury than young mice (**Supplementary Fig. 10**). Differential gene expression analyses showed that the mRNA levels of complement and coagulation cascade genes such as C4b (Adjust p1.66X10^-36^), C3 (Adjust p=1.14X10^-7^), as well as GFAP (Adjust p=0.003), were higher in the old ipsilateral hippocampi than in the young ipsilateral hippocampi. These data indicated that, like cortex, old mice also have a higher inflammatory response to stroke injury than young mice in the hippocampus.

### Transcriptional response to ischemic stroke injury in an age-dependent hemispheric pattern

WGCNA showed a marked difference in gene expression between the atrophic region and the corresponding contralateral cortex in old mice, suggesting an age-dependent transcriptional divergence in response to stroke. In young mice, gene expression patterns remain more consistent between the atrophic region and the corresponding contralateral cortex (**Fig. 8a & b**). This difference between the atrophic region and the corresponding contralateral cortex in old mice might reflect enhanced or altered neuroinflammatory and repair mechanisms triggered by ischemic injury, which appear to be less pronounced or differently regulated in young brains.

**Fig. 8.**
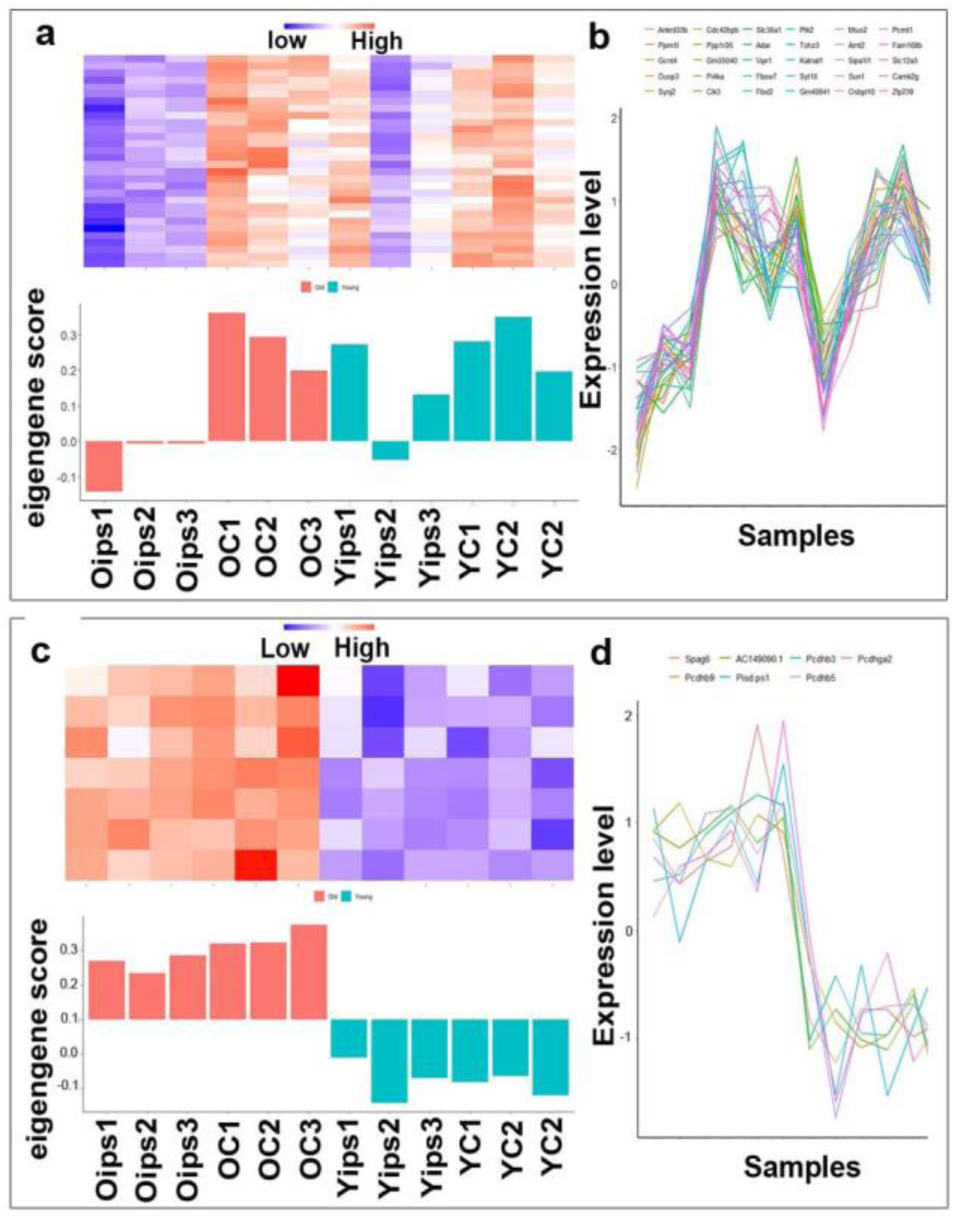
Age-dependent hemispheric transcriptional response to stroke injury **(a).** Heatmap (top). Relative gene expression levels across samples. Bar plot of eigengene scores (bottom) for a key module, showing significant differences between the atrophic region and the corresponding contralateral cortex in old mice but not in young mice. Oips: atrophic region of old mice; OC: corresponding contralateral cortex of old mice; Yips: atrophic region of young mice; YC: corresponding contralateral cortex of young mice. **(b).** Gene expression profiles. Line plot of individual gene expression within a module, in which each line represents a gene. The top 30 genes across all cortical samples exhibit a strong co-expression pattern. **(c).** Heatmap (top). Relative gene expression levels across hippocampal samples. Bar plot of eigengene scores (bottom) for a key module, showing significant differences between old and young mice. Oips: ipsilateral hippocampi of old mice; OC: contralateral hippocampi of old mice; Yips: ipsilateral hippocampi of young mice; YC: contralateral hippocampi of young mice. **(d).** Gene expression profiles. Line plot of individual gene expression within a module, in which each line represents a gene. The top 7 genes across all hippocampal samples exhibit a strong co-expression pattern.

We have also analyzed the hippocampal tissue from the same mice, focusing on age-related transcriptional responses in the hippocampi (**Fig. 8c & d**). Like the cortex, WGCNA revealed several gene modules with expression profiles that distinguished old mice from young mice. However, unlike the cortex, these hippocampal modules did not show significant differences between ipsilateral and contralateral sides within either age group. This lack of differentiation in the ipsilateral and contralateral hippocampi is consistent with its indirect exposure to the ischemic injury, as the stroke injury in our model directly affects the cortex, not the hippocampi.

### Activation of α7-nAchRs reduces the CD68^+^*/*SYN^+^ cells and GFAP^+^/SYN^+^ cells in the hippocampi

Transcriptome, protein, and histological analyses all show that old mice have more inflammatory cells, such as CD68^+^ and GFAP^+^, than young mice before and after stroke injury, which may cause excessive elimination of synapses and post-stroke memory dysfunction. We showed that tibia fracture 6 hours before ischemic stroke in young mice caused long-lasting post-stroke memory decline, which is associated with increased accumulation of activated microglia in the hippocampi [21, 28]. Alpha-7 (α-7) nAChRs are ligand-gated channels widely distributed on the surface of systemic macrophages [35] regulate inflammatory processes and signal through distinct intracellular pathways. The inflammatory process can be regulated by the activity of the parasympathetic nervous system, termed the cholinergic anti-inflammatory pathway [36]. Activation of α-7 nAchR has a neuroprotective effect on ischemic and hemorrhagic stroke [37]. We showed that activation of α7-nAchRs by PHA treatment reduced the activated microglia/macrophages in the hippocampi and improved the memory function of mice subject to tibia fracture 6 hours before ischemic stroke [28]. In this study, we found that old mice developed long-lasting memory dysfunction, which was also associated with more CD68^+^ activated microglia/macrophages and GFAP^+^ reactive astrocytes in the peri-atrophic regions and hippocampi than young stroke-only mice at 8 weeks after stroke injury (**Supplementary Figs. 7, 8, & 9**). More synapses were engulfed by CD68^+^ microglia/macrophages and GFAP^+^ reactive astrocytes in the peri-atrophic regions and hippocampi of old stroke mice than young stroke-only mice (**Fig. 2**). To test if activation of α7-nAchRs activity reduces synapses engulfing CD68^+^ microglia/macrophages and GFAP^+^ reactive astrocytes, we analyzed the number of CD68^+^ activated microglia/macrophages, GFAP^+^ reactive astrocytes, CD68^+^/SYN^+^ and GFAP^+^/SYN^+^ cells in the hippocampi of PHA treated young mice subjected to tibia fracture 6 hours before ischemic stroke at 8 weeks after the injuries. We found that PHA treatment reduced GFAP^+^ astrocytes (p=0.016) in the hippocampi ipsilateral to stroke injury but not CD68^+^ microglia/macrophages (**Supplementary Fig. 11**). PHA treatment also reduced CD68^+^/SYP^+^ on both sides of the hippocampi (p<0.001, **Fig. 9a & b**) and GFAP^+^/SYP^+^ cells in the hippocampi ipsilateral to stroke injury (p<0.001, **Fig. 9c & d**).

**Fig. 9.**
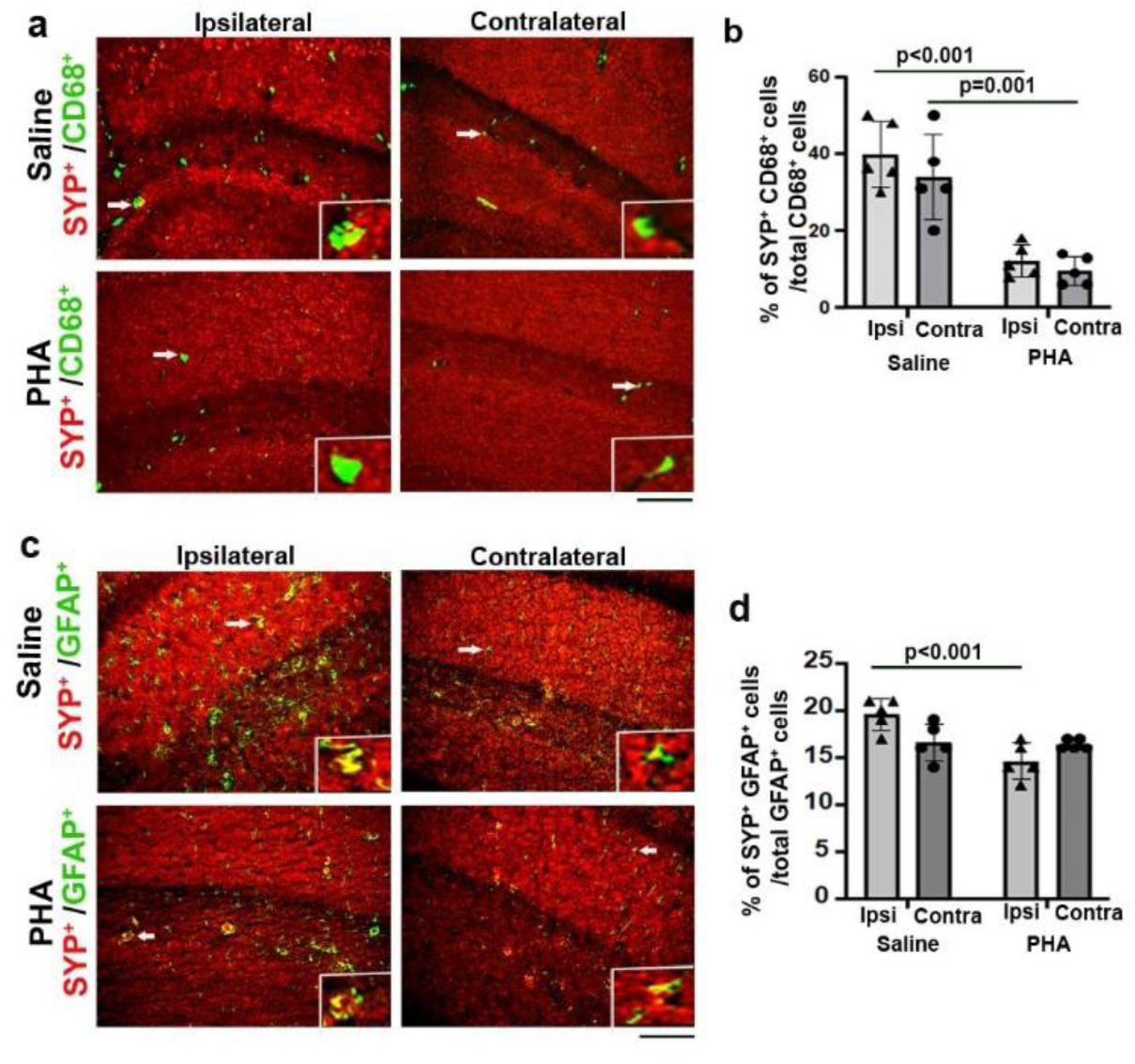
PHA treatment reduced CD68^+^/SYN^+^ microglia/macrophages and GFAP+/SYN^+^ astrocytes in the hippocampi ipsilateral to stroke injury of mice with tibia fracture shortly before stroke **(a).** Images of hippocampal regions co-stained with antibodies specific to CD68 (green) and SYN (red). Scale bar=50µm. The insertions in each picture show colonization of CD68 and SYN in the cells indicated by arrows. **(b).** Quantification of CD68^+^/SYN^+^ cells among total CD68^+^ cells. N=5. **(c).** Images of hippocampal regions co-stained with antibodies specific to GFAP (green) and SYN (red). Scale bar=50 µm. The insertions in each picture show colonization of GFAP and SYN in cells indicated by arrows. Scale bar=50µm. **(d).** Quantification of GFAP^+^/SYP^+^ cells among total GFAP^+^ cells. n=5. Ipsi: ipsilateral to stroke injury; Contra: contralateral to stroke injury.

These results demonstrated that activation of α7-nAchRs through PHA treatment reduces synapse removal by CD68^+^ microglia/macrophages and GFAP^+^ astrocytes in the hippocampi of mice subjected to tibia fracture 6 hours before ischemic injury. Therefore, inhibition of neuroinflammation by activation of α7-nAchRs can reduce synapse loss, which is one of the mechanisms of PHA-induced improvement of post-stroke cognitive function we have observed in a previous study in mice subjected to tibia fracture shortly before stroke injury [28]. These data support that excessive removal of synapses by inflammatory cells caused by enhanced neuroinflammation by comorbidities, such as bone fracture or aging, contributes to the long-lasting memory dysfunction post-stroke.

## Discussion

Stroke is a disease of aging, with approximately two-thirds of all strokes occurring in older adults [38]. Old mice exhibit a differential response to stroke and have worse outcomes than young mice [39]. In this study, we found that the old mice (15-18 months of age) developed long-lasting memory dysfunction (beyond 8 weeks) after ischemic stroke, a phenotype that is not observed in young mice with stroke injury only. The old mice had more activated microglia/macrophages (CD68^+^) and reactive astrocytes (GFAP^+^) in the peri-atrophic and hippocampal regions ipsilateral to stroke injuries than young stroke mice. There were more synapse-engulfing microglia/macrophages and astrocytes in the peri-atrophic region and hippocampi in the old mice than young mice, which was accompanied by a reduction of the complexity of neurons and dendritic spin densities in the cortex and hippocampi. The RNAseq data were consistent with the histological and protein analyses that the inflammatory pathways were upregulated after stroke in both young and old mice and were more robust in old mice. Pathways that regulate synapse organization, dendrite development, axonogenesis, axon development, dendrite morphogenesis, and neural differentiation were down-regulated in the ipsilateral cortex of old mice compared to young mice, suggesting more severe neuronal damage in the old mice after ischemic stroke. Pathways that regulate axonal part and axonogenesis, and dendrite membrane, were down-regulated in the ipsilateral hippocampi of old mice compared to the ipsilateral hippocampi of young mice and the contralateral hippocampi of old mice, as well as in the ipsilateral hippocampus of young mice compared to the contralateral hippocampi of young mice. WGCNA showed a marked difference in gene expression between the atrophic region and the corresponding contralateral cortex in old mice, suggesting an age-dependent transcriptional divergence in response to stroke. These data suggest a more severe neuronal damage in the old mice after ischemic stroke.

We showed that old mice, but not young mice, developed long-term memory dysfunction after ischemic stroke. Post-stroke memory dysfunction is devastating and common, and an underdiagnosed complication following stroke [40, 41]. Despite modest improvement in stroke outcomes [42], post-stroke memory dysfunction remains highly prevalent and disabling [43, 44]. Consistent with age-concordant ischemic injury [45, 46], we showed previously that old mice had more severe neuroinflammation and behavioral dysfunction at the acute stage of ischemic stroke [47]. We found, in this study, a stronger inflammatory response and oxidative stress in the peri-atrophic region in old mice than in the young mice at the chronic stage of ischemic stroke injury. This data indicates that ischemic stroke causes more severe neuroinflammation and neuronal injury in old mice, which may explain why elder patients have unfavorable outcomes after ischemic stroke than younger patients.

With aging, microglia acquires a dysfunctional phenotype characterized by dystrophic morphology, impaired phagocytosis, reduced motility, produce higher levels of reactive oxygen species, and have an exaggerated inflammatory response after ischemic stroke compared to young animals [48, 49]. Microglia can act indirectly as synaptic regulators without physical contact with neurons. When exposed to inflammation, microglia secrete extracellular vesicles that contain proteins, lipids, and RNAs. Internal miR-146a-5p targets and inhibits the expression of presynaptic synaptotagmin1 (Syt1), resulting in a reduction in the number of dendritic spines [50]. Neuronal exosomes promote microglial phagocytosis and enhance synaptic pruning. Incubation of microglia with neuron-secreted exosomes upregulated the expression of C3 in microglia and enhanced microglial clearance of inappropriate synapses [51]. Through RNAseq, we found significant down-regulation of Syt1 in the old periathrophic region compared to the contralateral part. A study showed that activation of microglia causes excessive loss of dendritic spines after transient global cerebral ischemia [52]. Therefore, the increase of activated microglia in the brain of old mice before and after ischemic stroke may contribute to the spin loss and long-lasting post-stroke memory dysfunction we have observed in this study.

A recent study identified a member of the C-type lectin receptor family, Clec7α, as a receptor involved in microglial phagocytosis. They found that Clec7α is required for synaptic engulfment by microglia after ischemic stroke. Clec7a has been shown to regulate phagocytosis by macrophages in multiple peripheral tissues. Moreover, Clec7α is upregulated in microglia during neurodegeneration and is an important receptor for microglial activation in response to AD pathology. Furthermore, Clec7α signals through spleen tyrosine kinase (SYK) to enhance the phagocytosis of A𝛽 [53]. Another study revealed the critical role of microglial Clec7α in long-term neurological deficits following ischemic stroke. Regulating the expression of microglial Clec7α may be an effective strategy for promoting long-term neurological function recovery after ischemic stroke [54]. Our RNAseq analysis shows that Clec7α gene expression is upregulated in the brains of old mice with or without stroke injury and in the hippocampi of old mice after stroke. The role of Clec7α in synaptic removal and long-lasting post-stroke memory dysfunction in old mice needs to be studied future.

Microglia and astrocytes change their features dramatically in response to stroke [55, 56]. Both microglia and astrocytes have enriched synapse engulfment pathway-related genes, including phagocytic receptors, intracellular molecules, and opsonins [57]. In the stroke brain, dead neurons and synaptic debris were found in microglia/macrophages and astrocytes [58, 59]. While microglia/macrophage and astrocyte-mediated phagocytosis are generally assumed to be necessary for clearing synaptic debris and beneficial for brain recovery, phagocytic microglia/macrophages and astrocytes also damage viable neurons in the stroke brain [60]. Although early studies have highlighted the important roles of microglia/macrophages and astrocytes in neuronal debris clearance during the early stage of stroke, Shi et al. showed that they are still phagocytic and cause synapse loss at the subacute stage (14 days after stroke) [19]. We analyzed whether microglia/macrophages and astrocytes phagocytose synapses at the chronic stage of ischemic stroke and if they contribute equally to synapse removal. We found that although more CD68^+^ microglia/macrophages and reactive astrocytes were engulfing synapses in old mice in the peri-atrophic region and hippocampi ipsilateral to stroke injury than in young mice at a later stage of ischemic stroke, 10-fold more GFAP^+^ cells were detected in old hippocampi, both ipsilateral and contralateral sides, than CD68^+^ cells. There were more GFAP^+^/SYN^+^ cells than CD68^+^/SYP^+^ cells in the peri-atrophic regions and in the ipsilateral hippocampi. These data indicate that reactive astrocytes contribute more than microglia to synapse removal at the chronic stage of ischemic stroke in old mice.

Astrocytes can act as antigen-presenting cells. The degradation of ingested materials has to be slow to present antigens to T-helper cells. Engulfed materials can be stored in astrocytes for an extended period before being degraded in lysosomes [61]. Lysosomal dysfunction and abnormal autophagosome accumulation have been noticed in aging astrocytes, which can lead to impaired synapse elimination [62, 63] and may cause more GFAP^+^/SYN^+^ cells than CD68^+^/SYP^+^ cells in old mice. However, many studies support that astrocytes play important roles in synapse removal in different conditions. Chung et al measured the phagocytic indices of astrocytes and microglia together in the dorsal lateral geniculate nucleus (LGN) during postnatal day 3 (P3)–P9 and found that between P3– P6, microglia engulfed more synapses than astrocytes per unit cell volume. Astrocytes significantly exceeded microglia by 10-, 7-, 6-and 4-fold at P5, P6, P7 and P9, respectively. Consequently, the total amount of engulfed synapses in a given imaging field was much greater in astrocytes during P3–P9, indicating that the total amount of synaptic pruning by astrocytes could outnumber that by microglia in the developing LGN [64]. Another study found that astrocytes play an important role in the elimination of excitatory and inhibitory synapses in the hippocampal CA1 region of the adult mouse [65].

It has been shown that the ApoD-induced inflammasome was activated in microglia, which gave rise to the proinflammatory phenotype. Targeting the microglial ApoD inhibited neuronal apoptosis [66]. These findings demonstrate the critical role of ApoD in microglial inflammasome activation. We found that ApoD is upregulated in the ipsilateral cortex and hippocampus of old mice compared to the young ipsilateral cortex and hippocampus and the old contralateral cortex following stroke. In addition to inducing an inflammasome in active microglia, ApoD also has a neuroprotective effect. Its increase in cognitive disorders could be a compensatory mechanism [66].

It has been shown that microglia are activated by complement component 3 (C3) released by astrocytes, causing microglia to consume and eliminate complement-tagged synapses [67]. The C3 bounds to synapses, triggering microglia synaptic engulfment [68]. Microglia and astrocytes have been reported as primary sources of C3. In the central nervous system (CNS), C3a-C3aR binding mediates synaptic plasticity and microglial phagocytosis. During the developmental period of synaptic refinement, astrocyte-secreted TGF-β increases C1q expression in neurons [69]. This results in the release of C3, which binds to C3R in microglia, promoting the engulfment of synapses. We found that C3 expression increased in the peri-atrophic region and ipsilateral hippocampus of old mice compared to the young peri-atrophic region and ipsilateral hippocampus at the chronic stage of ischemic stroke injury. C3 expression is also increased in the peri-atrophic region and ipsilateral hippocampus of old and young mice compared to the contralateral parts.

Astrocytes contact and eliminate synapses in a C1q-dependent manner [70]. C1q deletion reduced astrocyte–synapse association and decreased astrocytic and microglial synapse engulfment in Tau^P301S^ mice and rescued synapse density in AD patients [70]. We found up-regulation of complement and coagulation cascades in the uninjured cortex of old mice compared to young mice and the hippocampi of old stroke mice compared to young stroke mice, and a significant increase in C1q expression in the peri-atrophic region and ipsilateral hippocampi of old mice versus young peri-atrophic region and ipsilateral hippocampi at the chronic stage of ischemic stroke injury. C1q expression also increased in the peri-atrophic region and ipsilateral hippocampi of old and young mice compared to the corresponding contralateral parts. Unlike in disease conditions, synapse eating by astrocytes seems to be C1q-independent under physiological conditions [70]. Together, the above evidence indicates that complementary pathways could be a target for developing therapies to prevent post-stroke memory dysfunction.

We showed previously that young mice that underwent tibia fracture 6 hours before ischemic stroke developed long-lasting memory dysfunction, which coincided with an accumulation of CX3C chemokine receptor 1^+^ (Cx3cr1^+^) and CD68^+^ cells [21] and impairment of BBB in the hippocampi, and damage of the white matter in the striatum [22]. In contrast, ischemic stroke or tibia fracture alone causes temporary memory dysfunction (<1 week) in young mice [23, 71, 72]. Strong inflammatory response induced by ischemic stroke is a main pathological mechanism that contributes to stroke outcome. The cholinergic anti-inflammatory pathway inhibits cytokine release through activation of α7 nAChRs [73]. Treatment with an α7 nicotinic agonist, PHA, reduced neuroinflammation and post-stroke memory dysfunction in mice that had a tibia fracture 6 hours before ischemic stroke, partially through inhibiting cytokine release [28].

In this study, we found that synapse-engulfing microglia/macrophages and astrocytes were also increased in the ipsilateral hippocampi in young mice that underwent tibia fracture 6 hours before ischemic stroke. Our RNA-seq results indicated that the response to acetylcholine and cholinergic synaptic transmission was down-regulated in the old ipsilateral cortex compared to the young cortex after stroke. Nicotinic transmission in the hippocampus modulates cognitive function [74]. In a recent study, activating α7nAChR ameliorated cognitive dysfunction by upregulating the ERK1/2/CREB signaling pathway and reducing cell apoptosis in the hippocampal tissue of mice [75]. Therefore, we tested whether PHA treatment decreases the synapse removal in the hippocampi of mice that had a tibia fracture 6 hours before ischemic stroke. We found that PHA treatment reduced SYN-positive microglia/macrophages and astrocytes. These data suggest that excessive synapse removal by inflammatory cells is one of the underlying mechanisms contributing to long-lasting post-stroke memory dysfunction in stroke mice.

## Summary

In this paper, we showed that old mice developed long-lasting memory dysfunction (beyond 8 weeks) after ischemic stroke, a phenotype that had not been observed in young stroke only mice. Old mice had stronger pro-inflammation response to stroke injury and more synapse engulfing microglia/macrophages and astrocytes in the peri-atrophic region and hippocampi than young mice, which were associated with reduction of the complexity of neurons in the cortex and hippocampi. Therefore, enhanced neuroinflammation resulting in more synapse removal in old mice after ischemic stroke could be one of the mechanisms contributing to long-lasting post-stroke memory dysfunction in old mice. There were more synapse-engulfing astrocytes than synapse-engulfing microglia/macrophages in the peri-atrophic region and hippocampi, suggesting that astrocytes play a more important role in synapse removal at the chronic stage of ischemic stroke. We previously reported that the reduction of neuroinflammation by PHA treatment reduced post-stroke memory dysfunction in mice with tibia fracture shortly before ischemic injury [28]. In this study, we showed that PHA treatment reduced SYN-positive microglia/macrophages and astrocytes in the hippocampi of mice with tibia fracture shortly before ischemic injury. Therefore, excessive synapse removal by inflammatory cells may contribute to long-lasting post-stroke memory dysfunction of mice with comorbidities such as bone fracture and aging, which is consistent with a recent study, which showed that synaptic markers are associated with cognitive decline in AD patients [76]. Inhibition of synapse removal by inflammatory cells could be developed to reduce post-stroke memory dysfunction.

## Acknowledgment

We thank the members of the Center for Cerebrovascular Research for providing us constructive suggestions, Saeed Sadigh Eteghad for assisting behavioral test analysis, and Raaga Karumanchi for assisting RNAseq data analysis.

## Author contributions

H.S, Z.S, and Q.L conceived and designed the study, and P.P, Q.L, K.H performed animal modeling and brain sample collection. Z.S, Z.W, K.L, S.S, A.Y, J.S, C.W, and S.S.E contributed to data collection and data analysis, while G.C, J.H, A.F, provided expert advice about the study design and interpretation of the data. L.P. and P.H.J. provided critical input for the interpretation of data. Figures and first draft of the manuscript were prepared by Z.S, H.S, P.P. All authors provided feedback and approved the final version of the manuscript.

