## Supplementary material for "Increased Synapse Elimination by Inflammatory Cells Contributes to Long-lasting Post-Stroke Memory Dysfunction in Old Mice": Acta-supplementary file-8.docx

Center for Cerebrovascular Research

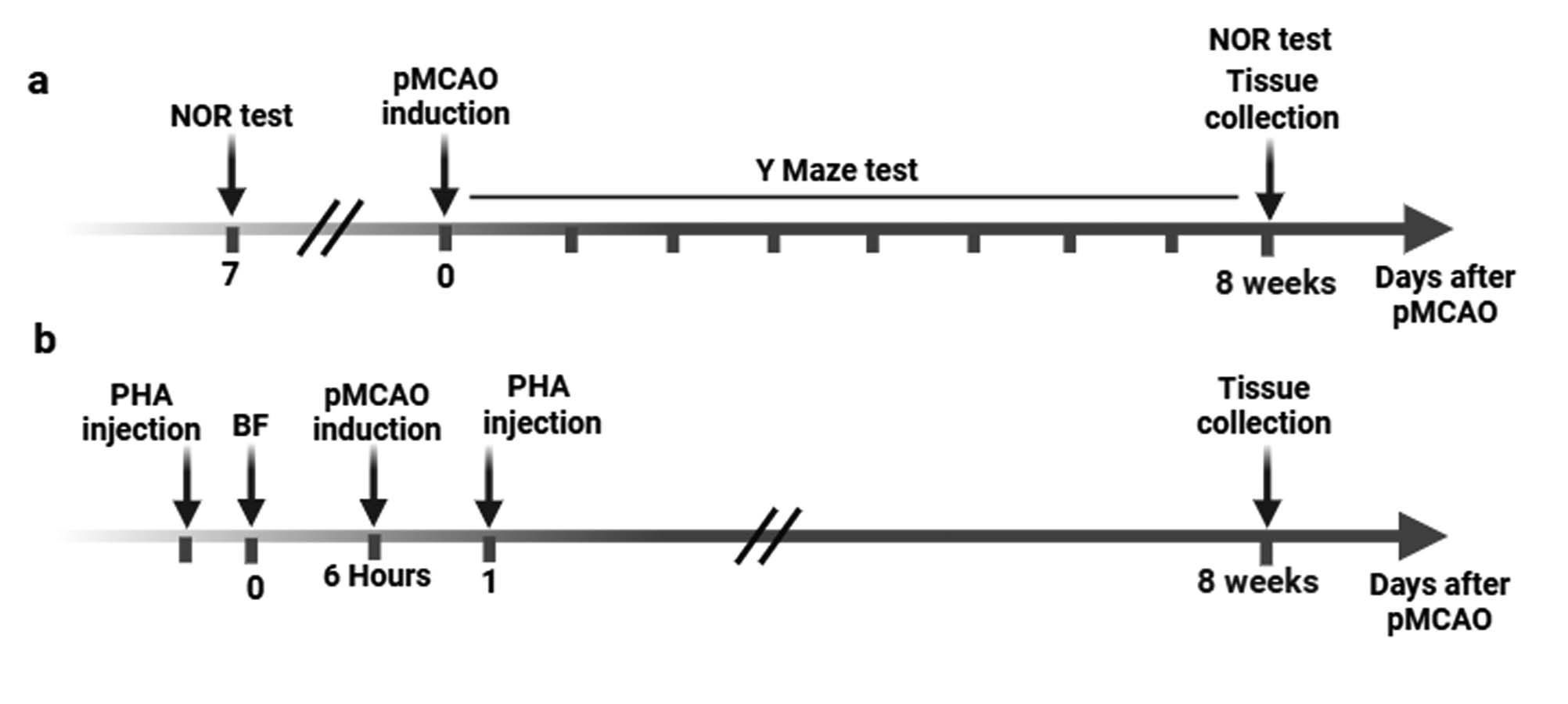

**Supplementary Fig. 1** Experimental design **(a).** pMCAO was induced in mice, the Y-maze test was performed every week, and the NOR test was performed 7 days before pMCAO and 8 weeks after pMCAO. Brain tissues were collected 8 weeks after the pMCAO. **(b).** PHA was given to mice right before the tibia fracture (BF) and one day after pMCAO. BF was performed 6 hours before pMCAO. Brain tissues were collected 8 weeks after pMCAO.

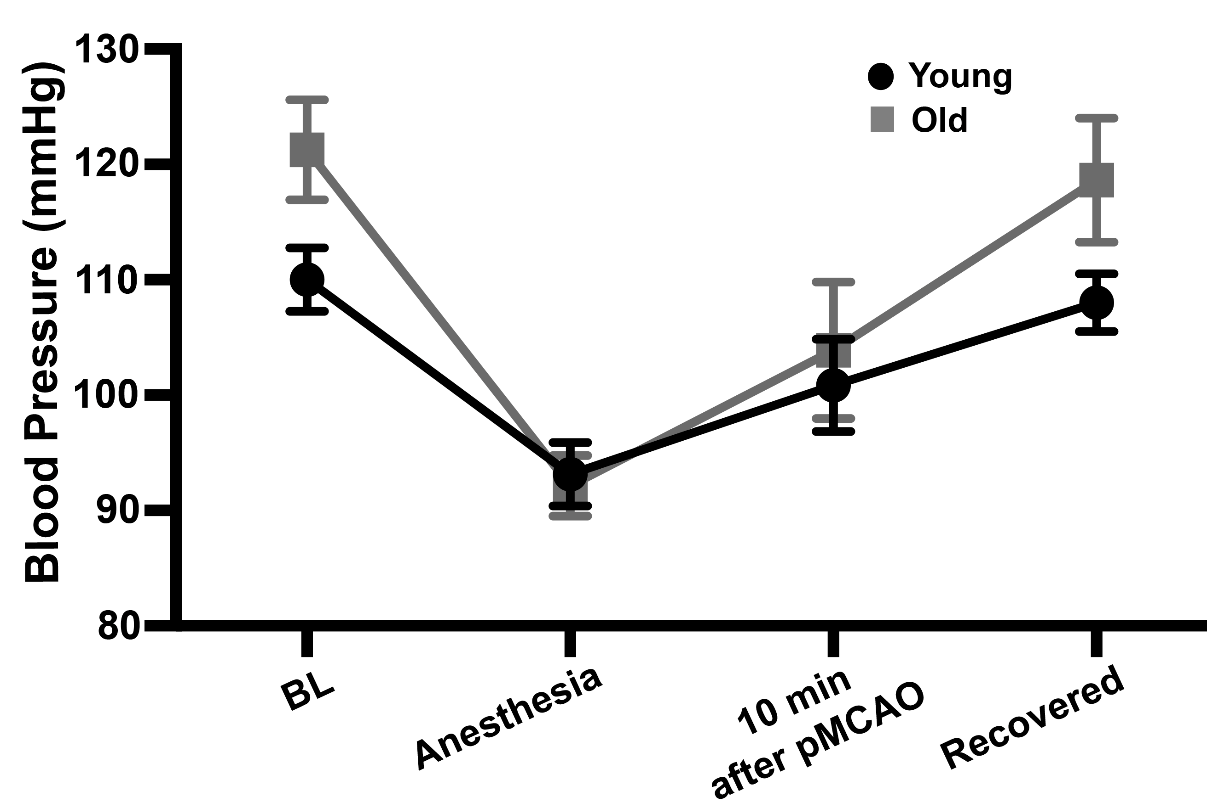

**Supplementary Fig. 2** Old mice had higher blood pressure at baseline (p<0.001) and the recovery stage of pMCAO procedure (p=0.001).

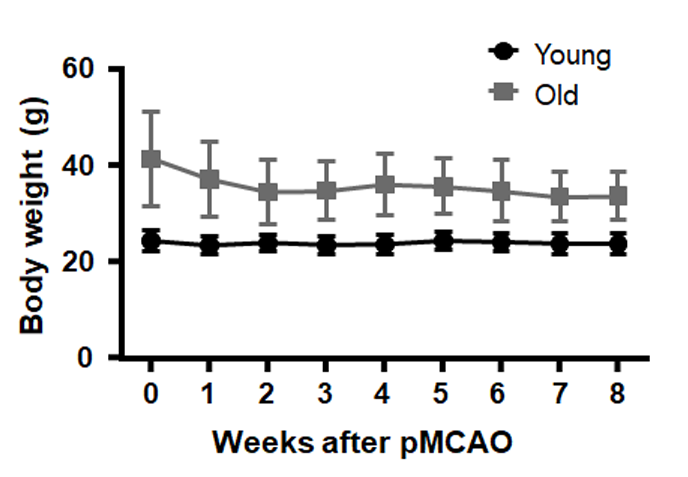

**Supplementary Fig. 3** The body weights of young and old mice were stable during the experiment, although old mice had higher body weights than young mice throughout the experimental period.

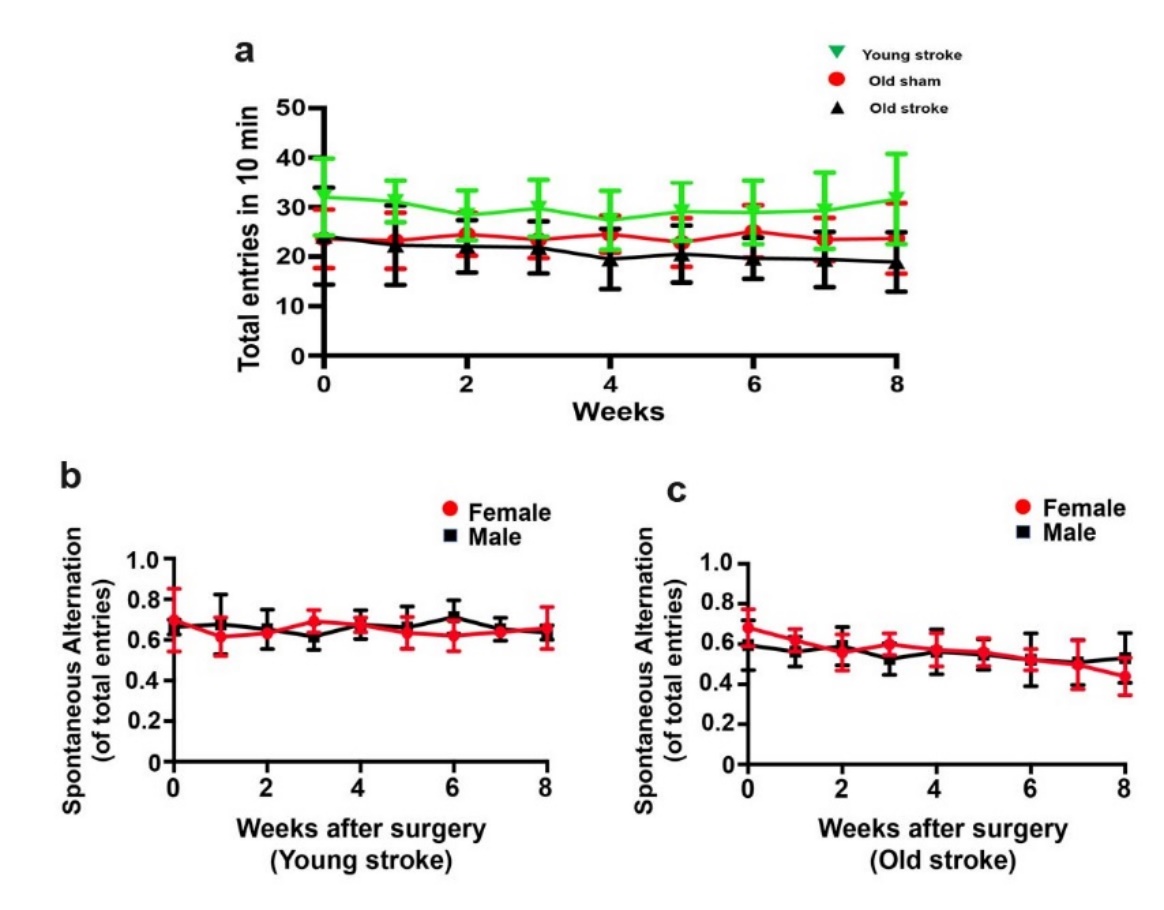

**Supplemental Fig. 4** Old mice with or without stroke made fewer entries in the 10-minute Y-maze test than young mice **(a).** Total entrances in 10 minutes. Young stroke mice made more entries than old stroke mice. Young stroke mice also made more entries than old sham-operated controls. n=15-16. **(b).** Young male (n=8) and female (n=7) mice made similar entries on Y-maze tests before and after stroke. **(c)**. Old male (n=8) and female (n=7) mice made similar entries on Y-maze tests before and after stroke.

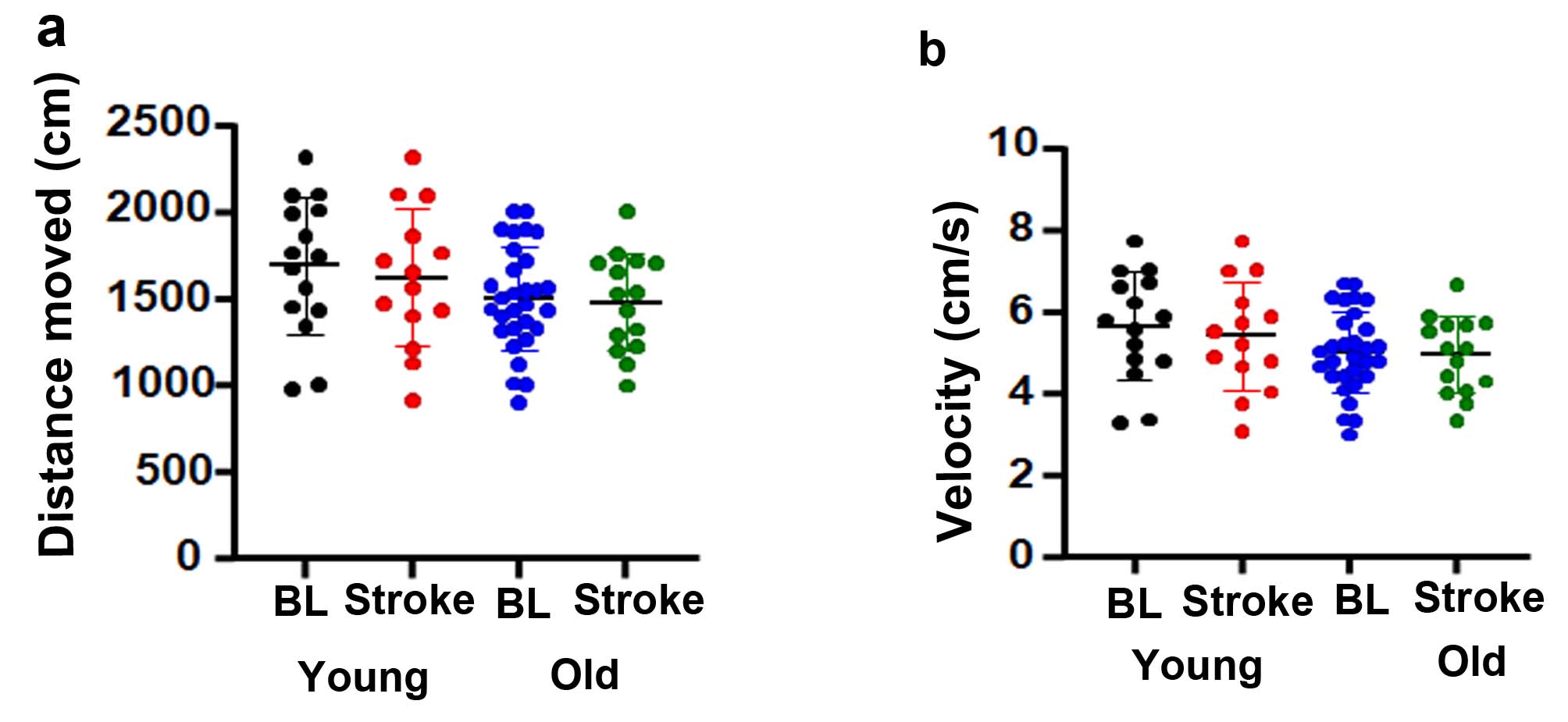

**Supplementary Fig. 5** The running distance and velocity are similar between young and old mice before and after stroke in the novel objective test **(a).** Running distance. **(b).** Running velocity**.** BL: baseline. N=15-30.

.

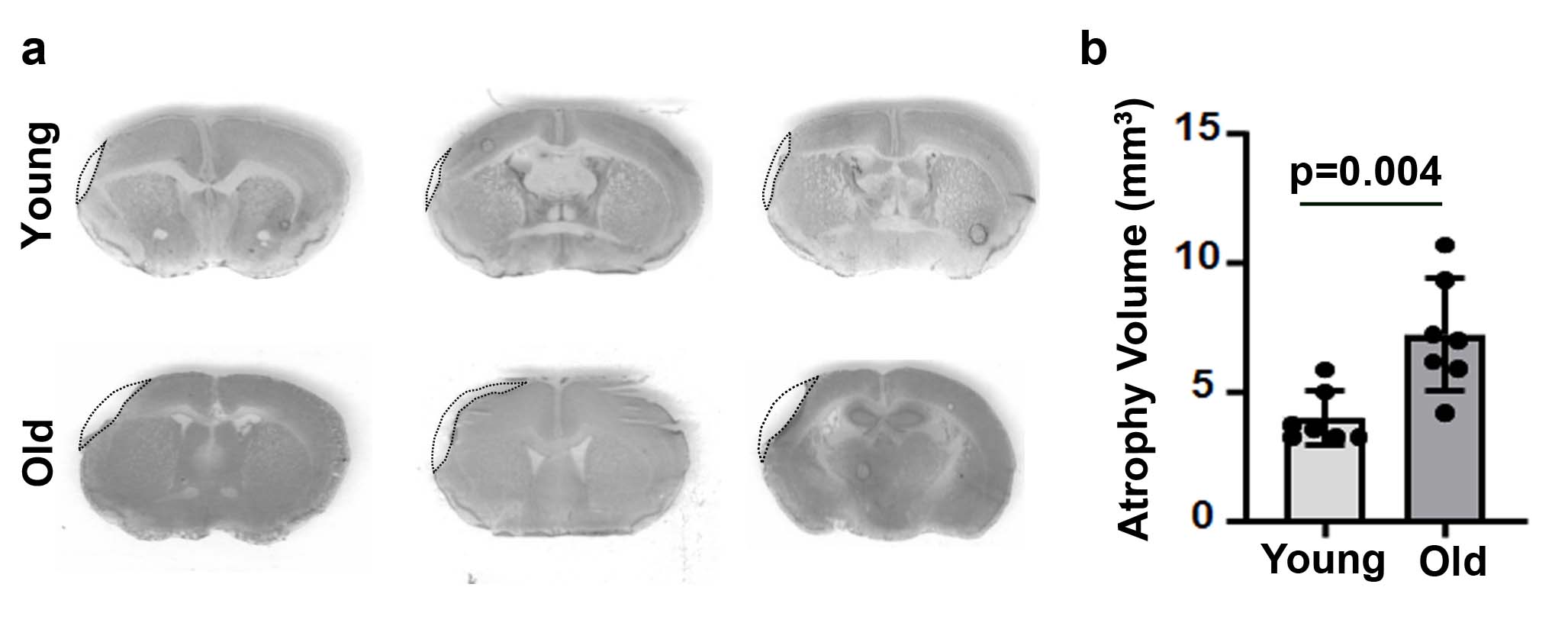

**Supplementary Fig. 6** Old mice have larger atrophic volumes than young mice 8 weeks after ischemic stroke **(a).** Images of cresyl violet-stained sections. **(b).** Quantification of atrophy volumes. N=7.

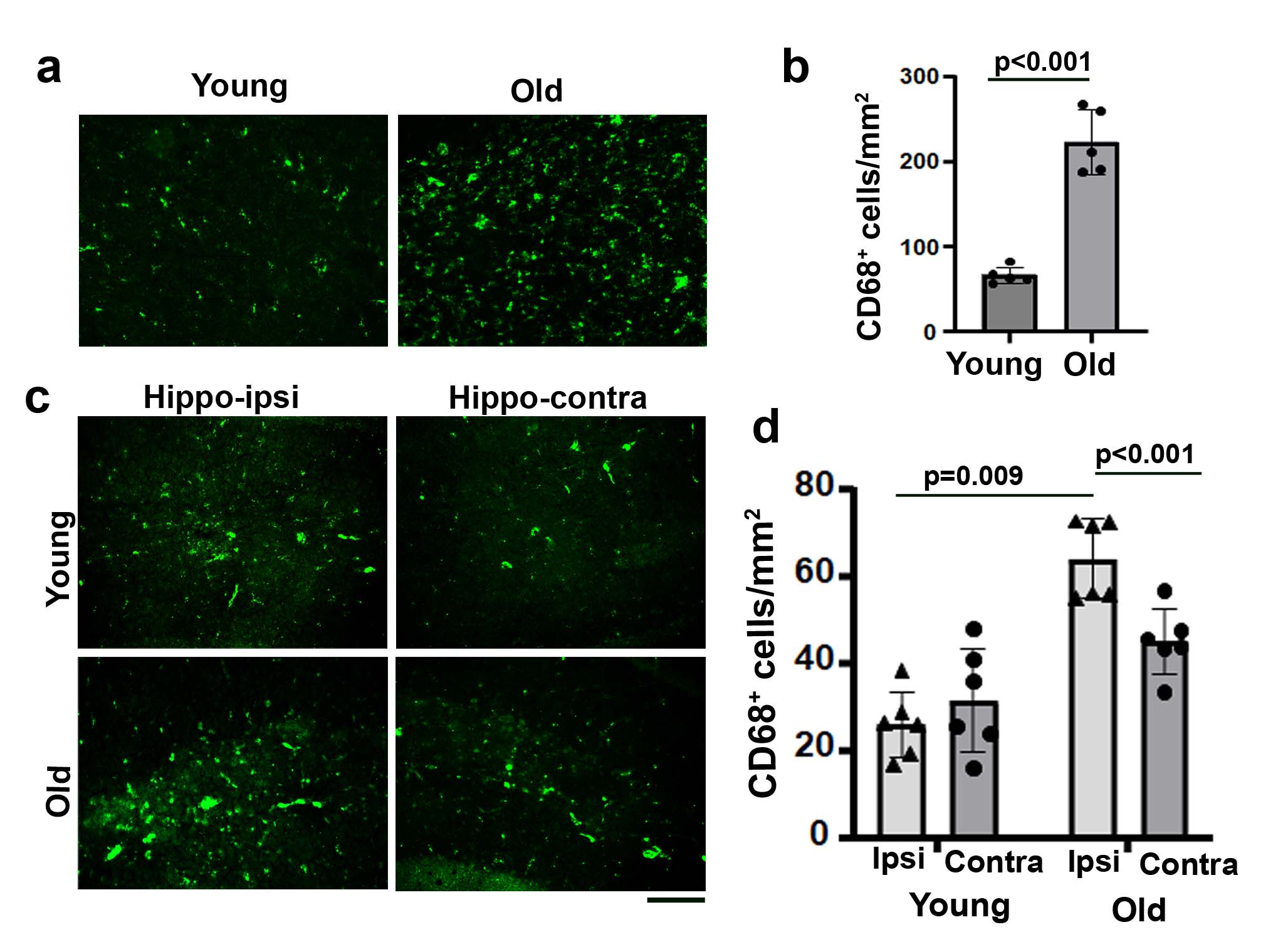

**Supplementary Fig. 7** Old stroke mice have more CD68^+^ cells in the peri-atrophic regions and hippocampi ipsilateral to stroke injuries than young stroke mice **(a).** Images of the peri-atrophic region stained with anti-CD68 antibody. CD68^+^ cells were stained green. **(b).** Quantification of CD68^+^ microglia/macrophages in the peri-atrophic region. N=5. **(c).** Images of hippocampi stained with anti-CD68 antibody (green). **(d).** Quantification of CD68^+^ microglia/macrophages in the hippocampi. N=6. Scale bar = 50 µm.

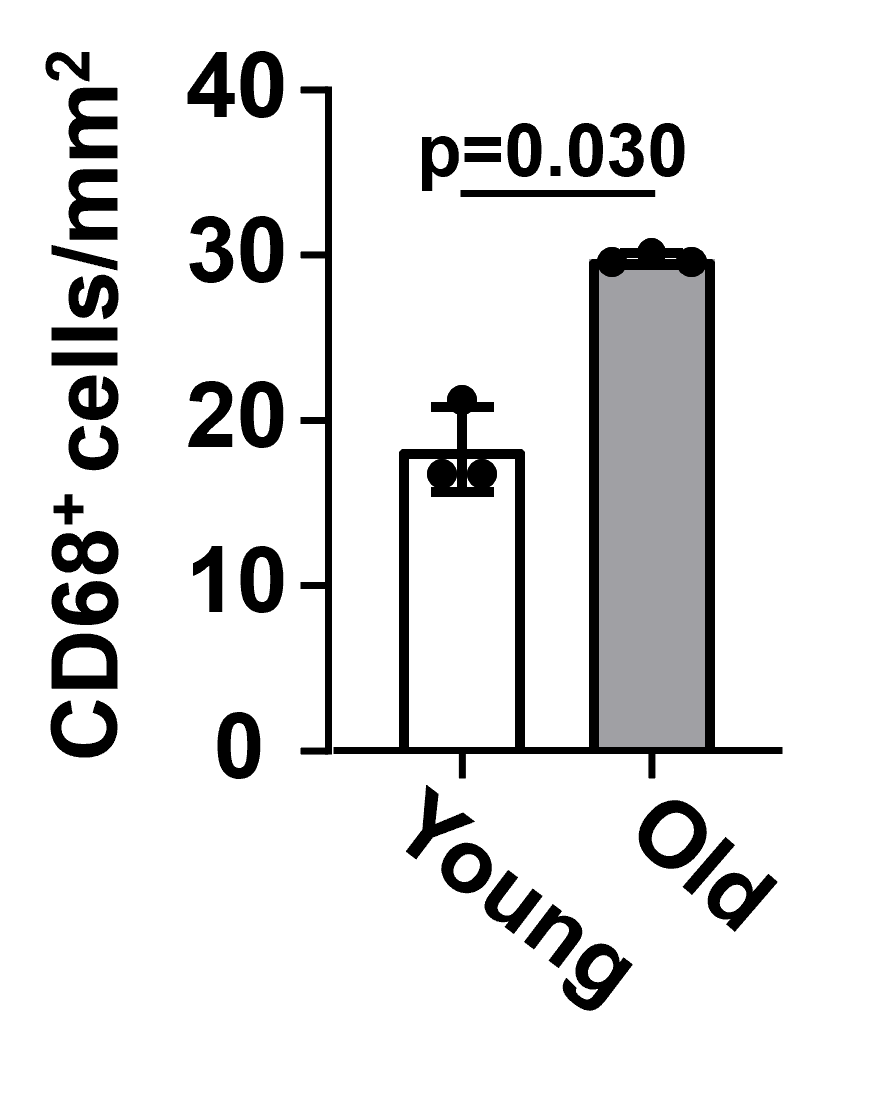

**Supplementary Fig. 8** Old mice have more CD68^+^ cells than young mice in the brain before stroke. N=3.

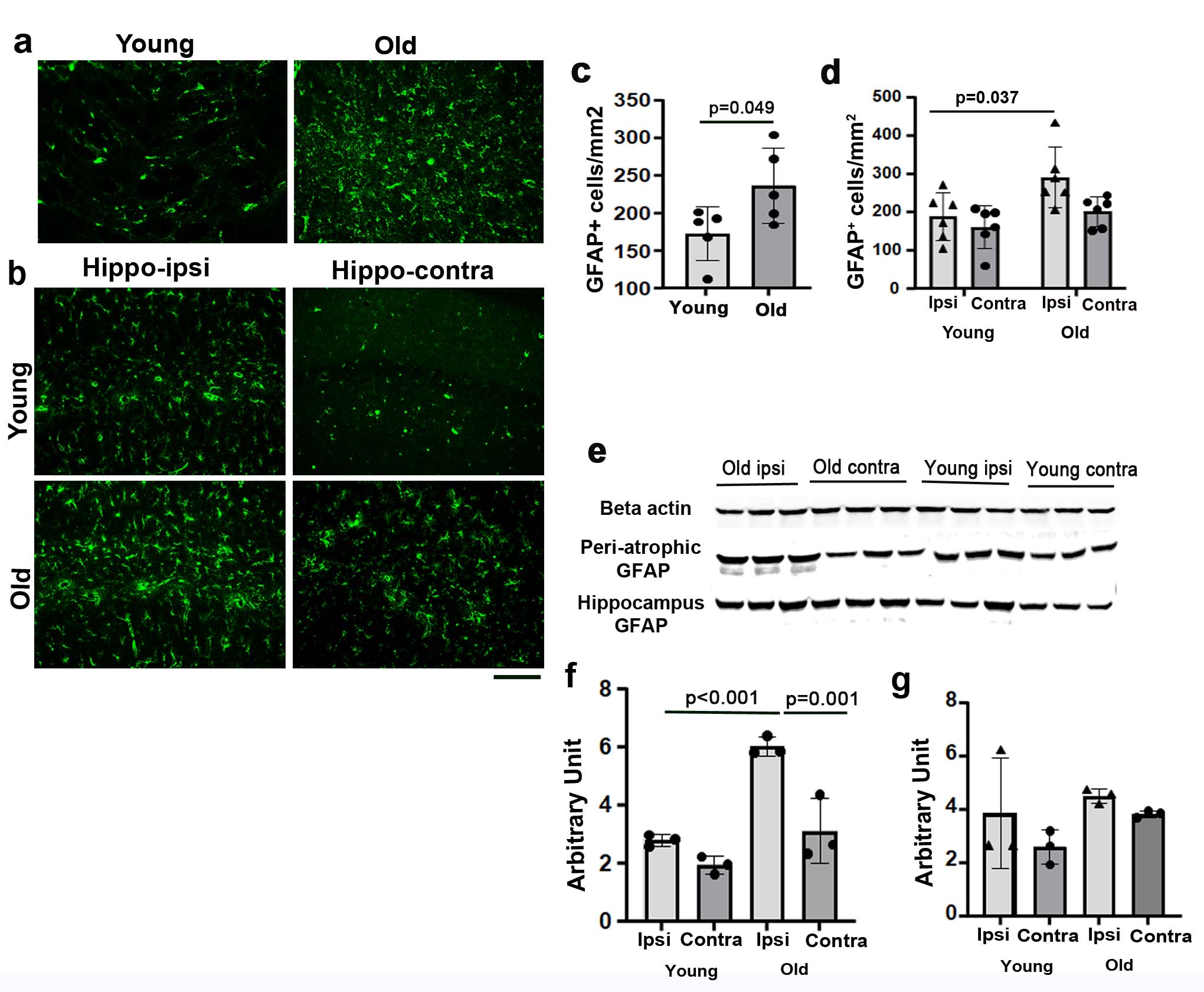

**Supplementary Fig. 9** Old stroke mice have more GFAP^+^ astrocytes in the peri-atrophic regions and hippocampi ipsilateral to stroke injuries than young stroke mice **(a).** Images of the peri-atrophic region stained with anti-GFAP antibody. GFAP^+^ cells were stained green. **(b).** Images of hippocampi stained with anti-GFAP antibody. Scale bar = 50 µm. **(c).** Quantification of GFAP^+^ cells in the peri-atrophic region. n=5. **(d).** Quantification of GFAP^+^ cells in the hippocampi. n=6. **(e).** Western blot images. **(f).** Quantification of GFAP protein in the atrophic regions and the corresponding contralateral uninjured cortex. n=3. **(g).** Quantification of GFAP protein in the hippocampi. n=3. Ipsi: ipsilateral to stroke injury; Contra: contralateral to stroke injury.

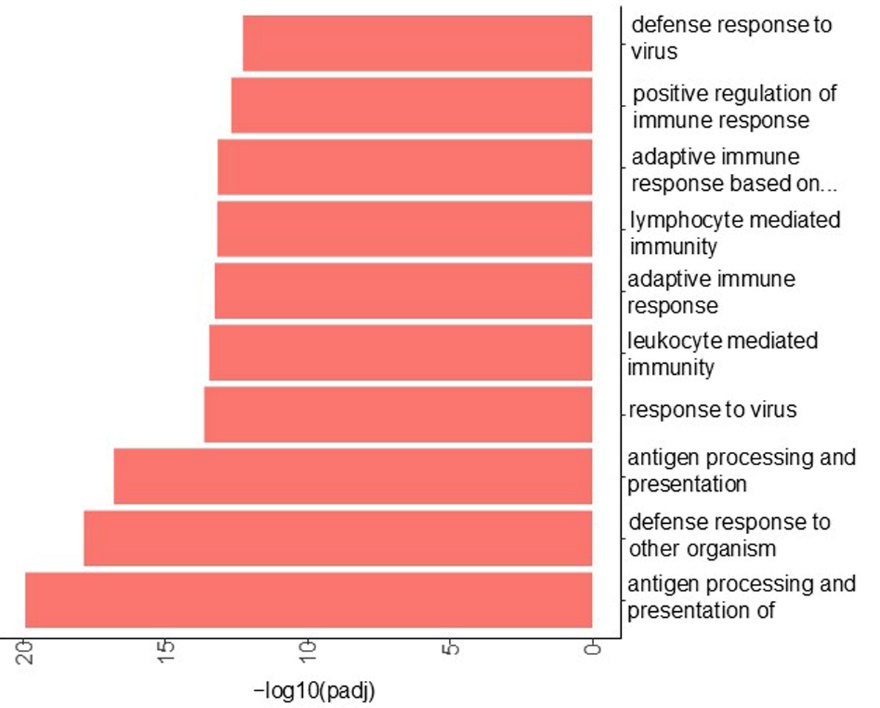

**Supplementary Fig. 10** Inflammation is upregulated in the ipsilateral hippocampi of old mice compared to young mice (top 10 upregulated biological pathways identified by GO analysis)

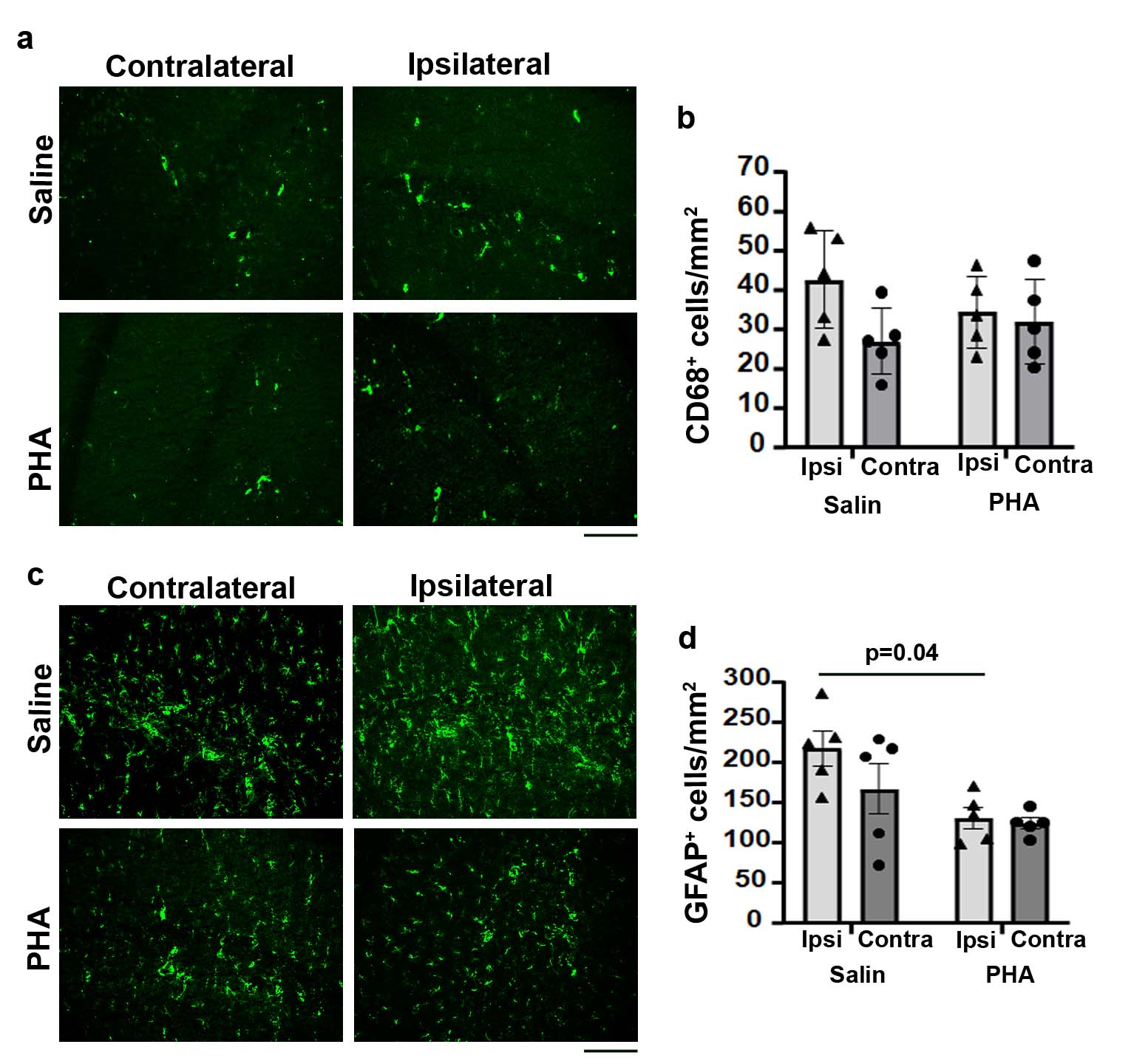

**Supplementary Fig. 11** Activation of α7-nAchRs by PHA treatment reduced the number of GFAP^+^ astrocytes in the hippocampi ipsilateral to stroke injury **(a).** Images of hippocampal sections stained with anti-CD68 antibody. CD68^+^ cells were stained green. Scale bar=50 μm. **(b).** Quantification of CD68^+^ microglia/macrophages in the hippocampi. n=6. **(c).** Images of hippocampal sections stained with anti-GFAP antibody. GFAP^+^ cells were stained green. Scale bar=50 µm. **(d).** Quantification of GFAP^+^ astrocytes in the hippocampal sections. n=5. Ipsi: ipsilateral to stroke injury; Contra: contralateral to stroke injury.
